# Extracellular protein catabolism drives regulated nitrogen handling and ammonia buffering in acute myeloid leukemia

**DOI:** 10.64898/2026.08.27.747222

**Authors:** Nina Kurrle, Philipp Makowka, Vera Schlipfenbacher, Johanna Kreitz, Islam Alshamleh, Marcel Seibert, Sifora Kaleab, Celestine Aguilar Montero, Silvia Marin, Dominik Fuhrmann, Florian Gatzke, Cláudia Fernandes, Nidhi Preman, Björn Häupl, Sebastian Wolf, Josefine Jakob, Amirhossein Barati Sedeh, Lea Fries, Leon Boerner, Heike Nuernberger, Verena Stolp, Rahul Kumar, Marlyn Thoelken, Christina Muhs, Christian Brandts, Janosch Martin, Martin Lindner, Tobias Berg, Jan Jacob Schuringa, Daniela S. Krause, Bernhard Brüne, Halvard Bonig, Sebastian Scheich, Marta Cascante, Thomas Oellerich, Harald Schwalbe, Frank Schnütgen, Hubert Serve

## Abstract

Acute myeloid leukemia (AML) cells exhibit pronounced metabolic plasticity, yet how amino acid supply is coordinated to sustain leukemic metabolism remains poorly understood. Here, we show that AML cells catabolize extracellular proteins as a major source of amino acids through lysosomal degradation of albumin. This proteocatabolic activity supports anabolic processes and mitochondrial energy production and establishes a regulated, high-throughput regime of primary nitrogen-containing metabolites (nitrogen regimen). Sustained proteocatabolism inevitably generates ammonia, and we find elevated ammonia concentrations in bone marrow plasma from newly diagnosed AML patients that decline with effective induction therapy. Using metabolomics, isotope tracing and targeted genetic and pharmacological manipulations, we identify glutamate-ammonia ligase (GS/*GLUL*) as a central enzyme that buffers proteocatabolism-derived ammonia by stabilizing intracellular nitrogen homeostasis. Loss of GS function limits sustainable nitrogen handling capacity, thereby impairing leukemic proliferation and delaying disease progression *in vivo*. Together, our findings define extracellular protein catabolism as a regulating nitrogen management strategy in AML and reveal GS as a capacity-defining vulnerability of proteocatabolic growth.

**Highlights:**

- AML cells utilize extracellular proteins as a major source of amino acids
- Proteocatabolism establishes a high-throughput nitrogen utilization regimen in AML
- Ammonia accumulation emerges as a predictable consequence of sustained proteocatabolism
- Glutamate-ammonia ligase (GS/*GLUL*) buffers ammonia and defines nitrogen handling capacity in AML

## 1. Introduction

Acute Myeloid Leukemia (AML) is an aggressive hematologic malignancy characterized by the clonal expansion of poorly differentiated myeloid blasts in the bone marrow (BM). Despite recent approval of new therapeutic agents and high initial remission rates following induction chemotherapy, relapse is common and for the vast majority of AML patients, allogeneic bone marrow transplantation is the only curative option. As a consequence, overall five-year survival rates for most AML patients are less than 30% despite aggressive therapies, highlighting the need to identify new, mechanism-based vulnerabilities of leukemic cells [1–3].

Metabolic reprogramming is a central feature of AML biology and contributes to leukemic growth, persistence, and therapy resistance [4–7]. Through clonal evolution and dynamic reprogramming during disease progression, AML cells exhibit remarkable metabolic plasticity that supports survival under therapeutic and microenvironmental stress. This flexibility enables AML cells to thrive in the BM microenvironment, where they disrupt normal hematopoiesis. As a result of this process, the metabolic wiring of AML cells is characterized by a high degree of cellular and patient-to-patient heterogeneity. However, multiple studies have identified critical dependencies on specific metabolites and pathways, including amino acid metabolism, redox regulation, and mitochondrial function [8–19]. In particular, several amino acids including glutamine (Gln), methionine (Met), cysteine (Cys), and branched-chain amino acids have been shown to be essential for AML survival and growth [10, 11, 13, 16, 17]. While these metabolic dependencies are well established at the level of individual nutrients and pathways, how AML cells coordinate amino acid supply and downstream nitrogen handling as an integrated metabolic program remains poorly understood.

One way by which cells can sustain their amino acid supply is through utilization of extracellular proteins as a nutrient source. Plasma proteins, with albumin as the most abundant circulating protein, represent a large and continuously available source of amino acids in serum [20]. In several solid tumors, uptake and lysosomal degradation of extracellular proteins enable cancer cells to convert this nutrient source into intracellular amino acids that support anabolic growth, mitochondrial metabolism, and signaling pathways, particularly under nutrient-limited conditions [21–27]. In these contexts, extracellular protein catabolism has largely been interpreted as an adaptive mechanism that supplements pools of free amino acids.

In this study, we show that AML cells catabolize extracellular proteins as a major source of amino acids. Leukemic blasts efficiently internalize and lysosomally degrade albumin, with particularly high activity in populations showing monocytic differentiation. This proteocatabolic activity supplies amino acid–derived nitrogen to central metabolic pathways, supporting anabolic processes and mitochondrial energy production within the BM microenvironment.

At high proteocatabolic throughput, amino acid catabolism inevitably generates ammonia as a biochemical byproduct that must be actively managed [28]. We find that extracellular protein catabolism in AML increases ammonia production *in vitro* and *ex vivo*, and that BM plasma from newly diagnosed AML patients contains elevated ammonia levels that decline upon effective treatment. These observations identify ammonia accumulation as a disease-linked feature that reflects ongoing nitrogen throughput rather than nonspecific toxicity.

Using metabolomics, isotope tracing, and functional perturbation approaches, we show that AML cells buffer these increased ammonia concentrations through coordinated activation of nitrogen-handling pathways [29, 30]. Central to this process is the enzyme glutamate–ammonia ligase (GS/*GLUL*), which incorporates excess ammonia into Gln and stabilizes intracellular nitrogen homeostasis [30, 31]). Loss of GS function limits the capacity of leukemic cells to tolerate proteocatabolic nitrogen flow, resulting in impaired proliferation and reduced leukemogenic potential *in vivo*.

Together, our findings define a metabolic framework in which extracellular protein catabolism enables AML cells to establish a regulated, high-throughput nitrogen regimen. Proteocatabolism supplies bulk reduced nitrogen that fuels anabolic and energetic demands, while ammonia accumulation emerges as a predictable spillover that constrains this process unless efficiently buffered. GS functions as a stabilizing node that sets the upper limit of sustainable nitrogen handling capacity, revealing an inherent and potentially exploitable vulnerability of proteocatabolic growth.

## 2. Materials and Methods

### 2.1. Cell culture

Human AML cell lines (HEL, MV411, MOLM-13, MOLM-14, THP-1, HL-60, PL-21, U937, ML-2, Kasumi-1, OCI-AML3, KG-1, NB-4, EOL-1, PLB985), human T-cell leukemia cell lines (JURKAT, MOLT-4) were obtained from the DSMZ (Deutsche Sammlung von Mikroorganismen und Zellkulturen GmbH). Human T-cell leukemia cell lines (CEM/C1, MOLT-3 and KE-37) and B-cell leukemia cell lines (RCH-ACV, SEM) were kindly provided by Dr. Anjali Cremer (University Medicine, Goethe University Frankfurt, Germany). Human AML cell lines stably expressing spCas9 were generated by lentiviral transduction (see 2.13) by using the plasmid LentiCas9-Blast (Addgene, #52962). Cells were cultured in RPMI 1640 medium (Gibco/ThermoFisher, #21875034) supplemented with 10% or 20% (for HT-93 and Kasumi-1) fetal calf serum (FCS, Sigma-Aldrich/Merck) and 100 IU/ml penicillin and 100 mg/ml streptomycin (P/S, Gibco/ThermoFisher, #15140122) at 37°C in a humidified 5% CO_2_ incubator. HEL cells overexpressing wildtype TFEB were generated by lentiviral transduction (see 2.13) as a mCherry-TFEB cDNA fusion. The TFEB WT cDNA was provided by Prof. Ritva Tikkanen (Justus-Liebig University Giessen, Germany). Lenti-X 293T cells (Takara, #Z2180N) were cultured in Dulbeccòs modified Eagle’s medium (DMEM 4.5 g/L D-glucose, Gibco/ThermoFisher, #41965-039) and supplemented with 10% FCS and 100 IU/ml penicillin and 100 mg/ml streptomycin (P/S, Gibco/ThermoFisher, #15140122) at 37 °C in a humidified 5% CO_2_ incubator.

To analyze the effect of glucose deprivation, cells were cultured in RPMI 1640 medium w/o glucose (#11879020, Gibco/ThermoFisher) supplemented with 10% fetal calf serum (FCS, Sigma-Aldrich/Merck), 100 IU/ml penicillin and 100 mg/ml streptomycin (P/S, #15140122, Gibco/ThermoFisher) and 0.5 g/L or 2 g/L glucose (D-glucose, #A2494001, Gibco/ThermoFisher). To analyze the effect of leucine or glutamine depletion, cells were cultured in RPMI 1640 medium without L-glutamine and L-leucine (#R8999-03, US Biological) prepared according to the manufacturer’s instructions. Medium was supplemented with the respective amino acid (0.38 mM L-leucine, #L8912, Sigma-Aldrich/Merck; 2.05 mM L-glutamine, #25030081, Gibco/ThermoFisher), fetal calf serum (FCS, Sigma-Aldrich/Merck) and 100 IU/ml penicillin and 100 mg/ml streptomycin (P/S, #15140122, Gibco/ThermoFisher) to obtain full medium, leucine- or glutamine-free medium. For bovine serum albumin addition, 5% (w/v) BSA (#A9418, Sigma-Aldrich/Merck) was added to the respective medium and the pH value was adjusted.

Human primary AML mononuclear cells were purified by Ficoll-Paque™ Premium gradient centrifugation according to the manufacturer’s instructions (GE Healthcare, USA). After purification, the cells were immediately stored in liquid nitrogen. The patients approved the sample collection and gave their consent according to the Declaration of Helsinki. The Ethics Committee of Frankfurt University Hospital also gave approval for the use of BM aspirates (approval no. SHN-11-2016/SHN-09-2019/SHN-7-2023). For experimental use, primary AML cells were defrosted and cultured in X-vivo 10 medium (Lonza, Switzerland) supplemented with 10 % (v/v) HyClone FCS (GE Healthcare, USA), 25 ng/ml hTPO (#300-18, Peprotech), 50 ng/ml hSCF (#300-07, Peprotech), 20 ng/ml hFLT3-Ligand (#300-19, Peprotech), 20 ng/ml hIL3 (#200-03, Peprotech), 100 IU/ml penicillin and 100 mg/ml streptomycin and 2 mM glutamine (GlutaMAX supplement, Gibco/ThermoFisher, #35050061). To simulate physiologic hypoxic conditions, primary AML cells were defrosted and cultured in a hypoxia chamber (Biospherix, X3 Xvivo System). Cells were kept at 37°C under 1% O_2_, 5% CO_2_ and 94% N_2_. This condition will further be termed hypoxia. Liquids used in the chamber, e.g. 1x phosphate buffered saline (1x PBS) and X-vivo medium were adapted to hypoxic condition over night for 12 hours.

Mobilized CD34^+^ hematopoietic stem/progenitor cells were isolated from left-over quality control material of mononuclear cell apheresis products of healthy adult volunteer donors undergoing G-CSF-mobilized stem/progenitor cell donation for matched-unrelated stem cell donation at the Department of Cellular Therapeutics and Cell Processing of the German Red Cross Blood Service (Frankfurt, Germany). Donor eligibility assessment and clearance was performed in agreement with national and international guidelines, as previously reported [32]. Stem/progenitor cell mobilization was induced with split-dose G-CSF of 7.5-10 µg/kg body weight*day for a total of nine doses before mobilization, following established protocols [33]. Samples were anonymized to the investigators; use of anonymized residual apheresis material was permitted by Ethics vote #329/10 with written informed consent of the donor. For enrichment, a CD34 MicroBead Kit (human, #130-046-702, Milteny Biotec) was used according to the manufacturer’s instructions. Prior to subsequent analysis, cells were cultured for two hours in X-vivo 10 medium (Lonza, Switzerland) supplemented with 10 % (v/v) HyClone FCS (GE Healthcare, USA), 25 ng/ml hTPO (#300-18, Peprotech), 50 ng/ml hSCF (#300-07, Peprotech), 20 ng/ml hFLT3-Ligand (#300-19, Peprotech), 20 ng/ml hIL3 (#200-03, Peprotech), 100 IU/ml penicillin and 100 mg/ml streptomycin and 2 mM glutamine (GlutaMAX supplement, Gibco/ThermoFisher, #35050061) before being analyzed.

### 2.2. Antibodies

Rabbit monoclonal antibody against human and mouse glutamate-ammonia ligase (GS) was purchased from Abcam (#ab178422). Rabbit monoclonal antibodies against LAMTOR1/C11orf59 (#8975S) eukaryotic translation initiation factor 4E binding protein 1 (4EBP1, #9644), phospho-4EBP1 Ser65 (#9451), ribosomal protein S6 kinase beta-1 (P70S6K, #34475), phospho-P70S6K Thr389 (#97596), transcription factor EB (TFEB, #37785), phospho-TFEB Ser211 (#37681), phospho-TFEB Ser122 (#87932), LAMTOR1 (#87786), Histone H3 (#9715) and β-Tubulin (#2146) were purchased from Cell Signaling. A mouse monoclonal antibody against SREBP2 (#557037) was purchased from BD Biosciences, an antibody against Vinculin (#V9264) was purchased from Sigma-Aldrich/Merck, an antibody against glyceraldehyde-3-phosphate dehydrogenase (GAPDH, #ab8245) was purchased from Abcam and a polyclonal rabbit antibody against SQLE was purchased from Proteintech (#12544-1-AP). The primary antibodies used for Western blotting were detected with an HRP-conjugated goat anti-rabbit (Dako) or anti-mouse (LI-COR Biosciences) antibody. For immunofluorescence the following secondary antibody was used: Goat-anti-rabbit IgG (H+L) Highly cross-adsorbed secondary antibody, Alexa Fluor 647 (#A-21244, ThermoFisher). Antibodies used in flow cytometric analysis are listed in table 1.

**Table 1:** List of primary antibodies used for flow cytometry.

| Antibody | Clone | Fluorochrome | Source |
| --- | --- | --- | --- |
| Mouse anti-human CD33 | P67.6 | APC | #340474, BD Biosciences |
| Mouse anti-human CD45 | HI30 | PE | #555483, BD Biosciences |
| Mouse anti-human IgG | G18-145 | PE | #555787, BD Biosciences |
| Rat anti-mouse CD117 | 2B8 | BV421 | #562609, BD Biosciences |
| Rat anti-mouse CD11b | M1/70 | PE | #553311, BD Biosciences |
| Rat anti-mouse Ly-6G (GR1) | 1A8 | APC | #560599, BD Biosciences |
| Mouse lineage antibody cocktail |  | APC | #558074, BD Biosciences |
| Rat anti-mouse Ly-6A/E (Sca1) | D7 | PE-Cy7 | #558162, BD Biosciences |
| Rat anti-mouse CD16/32 | 2.4G2 | PerCP-Cy5.5 | #560540, BD Biosciences |
| Rat anti-mouse CD34 | RAM34 | PE | #551387, BD Biosciences |
| CD34 human | 8G12 | FITC | #345801, BD Biosciences |
| Mouse anti-human CD45 | 2D1 | V450 | #642275, BD Biosciences |
| Mouse anti-human CD38 | HB7 | R718 | #567986, BD Biosciences |
| CD33 human | P67.6 | APC | #345800, BD Biosciences |
| Anti-human CD117 (c-kit) |  | BV785 | #313238, BioLegend |

### 2.3. Reagents

Gln solution (GlutaMAX Supplement, #35050061), DQ Red BSA (#D12051), Dextran Oregon green 488, 70,000MW (#D7172) were from ThermoFisher. Ammonium chloride (NH_4_Cl) (#A9434), Ammonium-^15^N-chloride (^15^NH_4_Cl) (#299251) and L-Methionine sulfoximine (#M5379) were obtained from Sigma-Aldrich/Merck. ^15^N-Arthrospira maxima hydrolysate was as kind gift of Silantes GmbH (Munich, Germany).

### 2.4 DQ-Red bovine serum albumin (BSA) protein catabolism assay

Cells lines (0.5*10^6^ cells/mL) were cultured for 24 hours in RPMI 1640GlutaMAX medium (#61870036, ThermoFisher) supplemented with 10% fetal calf serum (FCS, Sigma-Aldrich/Merck) and 100 IU/ml penicillin and 100 mg/ml streptomycin (P/S, #15140122, Gibco/ThermoFisher) at 37°C in a humidified 5% CO_2_ incubator. Primary AML samples were defrosted and cultured for 24 hours as described in 2.1. For the assay, cells were counted and resuspended in fresh medium in a final concentration of 1*10^6^ cells/mL. 2*10^5^ cells (cell lines) and 1*10^6^ cells (primary AML cells, CD34^+^ healthy donor cells) were treated for the indicated time points with DQ-Red BSA (stock concentration: 1 mg/mL; final concentration: 10 µg/mL) obtained from ThermoFisher (#D12051). After treatment cells were washed with ice-cold 1xPBS (#10010023, Gibco/ThermoFisher). Primary AML samples were stained at 4°C for 20 minutes with the surface antigen antibodies listed in section 2.2 (Table 1). Cells were analyzed by flow cytometry using a BD LRS Fortessa device (Becton Dickinson). Data were analyzed with the BD FACSDiva software (Becton Dickinson). For DQ-Red BSA filter G-610/20 was used.

To determine the intrinsic fluorescence of DQ-Red BSA, we measured DQ-Red BSA diluted in PBS in the absence of cells. DQ-Red BSA was prepared at a final concentration of 10 µg/mL by mixing 1 µL of a 1 mg/mL stock with 99 µL of 1× PBS in a black, flat-bottom 96-well plate (#655076, Greiner Bio-One). As a background control, 100 µL of 1× PBS was added to matched wells. Fluorescence was recorded using a Tecan Spark multimode plate reader in top-read mode with the following settings: excitation 590 nm (10 nm bandwidth), emission 620 nm (10 nm bandwidth), 10 flashes per well, 40 µs integration time, and a manually set Z-position of 20,000 µm. Gain was set automatically by the instrument (All measurements were performed at room temperature. Autofluorescence intensity of the DQ-Red BSA sample was background-subtracted using the PBS control.

To assess nonspecific binding and background fluorescence of DQ-Red BSA, we used both viable and fixed HEL cells. For fixation, HEL cells were incubated in 4% formaldehyde (P733.1, Carl Roth) for 20 min at RT, followed by two washes with 1× PBS prior to staining in RPMI 1640GlutaMAX medium as described above.

### 2.5 Dextran (70,000 MW) macropinocytosis assay

Cells lines (0.5*10^6^ cells/mL) were cultured for 24 hours in RPMI 1640GlutaMAX medium (#61870036, ThermoFisher) supplemented with 10% fetal calf serum (FCS, Sigma-Aldrich/Merck) and 100 IU/ml penicillin and 100 mg/ml streptomycin (P/S, Gibco/ThermoFisher, #15140122) at 37°C in a humidified 5% CO_2_ incubator. For the assay, cells were counted and resuspended in fresh medium in a final concentration of 1*10^6^ cells/mL. 2*10^5^ cells each were treated for the indicated time points with Dextran Oregon Green 488 (stock concentration: 2 mg/mL; final concentration: 0.05 µg/mL) obtained from ThermoFisher (#D7173). After treatment cells were washed with ice-cold 1xPBS (#10010023, Gibco/ThermoFisher). Data were analyzed with the BD FACSDiva software (Becton Dickinson). For Dextran Oregon Green 488 detection filter B-530/30 was used.

### 2.6 Immunfluorescence and confocal microscopy

Prior DQ-Red BSA (5 hours, see *2.4*) and Dextran staining (6 hours, see *2.5*) as well as immunofluorescent staining primary AML cells were sorted for CD45^+^/CD33^+^ and CD45^+^/CD34^+^ using a BD FACSAria III cell sorter, washed and transferred into a hypoxia chamber in X-vivo medium as described above (see *2.1*) and cultured for 1 hour.

HEL cells and primary AML cells (CD34^+^ and CD33^+^) were attached to microscope slides (Adhesive slides Superfrost Plus, #H867.1, Carl Roth) by centrifugation (Cytospin 4, Thermo Scientific) for 5 minutes and 500 rpm. Cells were fixed with 4% formaldehyde (37%, #P733.1, Carl Roth) for 10 minutes at room temperature (RT) followed by a concurrent permeabilization and blocking step with 1% bovine serum albumin (BSA) (w/v), 50 µg/ml digitonin (#D141, Merck) in 1x PBS for 30 minutes at RT. Thereafter, the cells were incubated with the primary antibody in 1% BSA (w/v) in 1x PBS for 1 hour, washed, incubated with the secondary antibody and 1 µg/mL 4’,6-diamidino-2-phenylindole (DAPI, #D9542, Sigma-Aldrich/Merck) for 1 hour and mounted in Fluoroshield with 1,4-Diazabicyclo[2.2.2]octane mounting medium (#F6937, Sigma-Aldrich/Merck). The samples were analyzed with Zeiss LSM 800 Confocal Laser Scanning Microscopes (Carl Zeiss).

### 2.7 DQ-Red BSA and comparison with routine diagnostic immunophenotyping

Flow cytometric data of DQ-Red BSA–treated primary AML samples were analyzed using FlowJo software (version 7.6.5; Tree Star Inc.). Viable cells were identified based on forward and side scatter properties and displayed in CD45 versus SSC-A plots to delineate blast and lymphocyte populations. For assessment of DQ-Red BSA uptake, gated populations were analyzed by plotting DQ-Red BSA fluorescence intensity (Comp-G-610-20) against SSC-A. Based on fluorescence intensity, cells were categorized as DQ-BSA^high, DQ-BSA^int, or DQ-BSA^low. These populations were backgated into the CD45/SSC-A plot and color-coded for visualization. Within the DQ-BSA^low fraction, lymphocytes were further distinguished from blast cells based on their CD45/SSC-A characteristics.

DQ-Red BSA–based flow cytometric analyses were compared with routine diagnostic immunophenotyping data obtained at the Department of Hematology, Medical Clinic II, University Medicine Frankfurt. Routine diagnostics were performed on freshly washed bone marrow samples without prior Ficoll density gradient separation using a BD FACSLyric™ flow cytometer (Becton Dickinson). Only antibodies that were included in the present analysis are reported. Routine data were reanalyzed using FlowJo software. Blast populations were identified in CD45/SSC-A plots and further analyzed for expression of CD117 (Comp-PE-Cy7-A) in combination with CD64 (Comp-PE-A) and CD11b (Comp-APC-A). CD11b- and/or CD64-positive and CD117-negative cells were classified as monocytoid-differentiated, whereas CD117-positive and CD11b/CD64-negative cells were classified as immature. These populations were color-coded and backgated into the CD45/SSC-A plot. Comparison of backgated populations revealed that DQ-BSA^high cells consistently displayed higher CD45 expression and SSC-A values than DQ-BSA^low cells and corresponded phenotypically to monocytoid-differentiated populations in routine diagnostic immunophenotyping. Patient samples included in the comparative analysis are summarized in Supplementary Table S1 (patient data used in the DQ-BSA assay).

### 2.8 Viability assay and IC_50_ determination

Viability/IC_50_ of EIPA, MSO, NH_4_Cl was assessed by measuring the ATP content by using the CellTiter-Glo luminescent cell viability assay (Promega, #G7570) according to the manufacturer’s instructions in AML cell lines and primary AML cells for 48 hours. Luminescence was measured in white Greiner 96 flat bottom polystyrene plates (#655074) using a Tecan infinite200Pro or Tecan Spark multimode plate reader.

### 2.9 Quantification of cell proliferation: cumulative growth assay and competitive growth assay

Cumulative growth assays were performed at a density of 0.25*10^6^ cells/ml if not otherwise stated, counted every 48 or 72 hours and then reseeded to the original density.

Competitive growth assays were performed by mixing EGFP-positive transduced cells at a ratio of (1:1) with non-transduced control cells that were monitored by flow cytometry every 48 hours using a BD LRS Fortessa device (Becton Dickinson). Data were analyzed with the BD FACSDiva software (Becton Dickinson).

### 2.10 Cell lysis, gel electrophoresis and Western blot

Cell pellets were lysed in SDS-lysis buffer (100 mM Tris-HCl pH 8, 150 mM NaCl, 10 mM EDTA, 10% SDS (w/v)) or RIPA-lysis buffer (50 mM Tris-HCl pH 7.5, 150 mM NaCl, 1% TritonX-100, 0.5% sodium desoxycholate, 0.1% SDS (w/v)) freshly supplemented with protease inhibitors (cOmplete™, Mini, EDTA-free Protease Inhibitor Cocktail, Roche/Merck, #11836170001). Lysates were cleared by centrifugation. Protein concentration was measured with the Bio-Rad DC Protein assay reagent (Bio-Rad, #5000111) or ROTI-Quant (Carl Roth, #K015.1), respectively. Equal protein amounts of the lysates were analyzed by SDS-PAGE and Western blot.

At least three independent experiments were performed for each assay, unless otherwise stated. For the quantification of Western blot protein bands, Image Studio 3.1 software (LI-COR Biosciences) was used. Data are shown as the mean ± SEM as stated in the figure legends. The different methods for statistical comparisons were applied using GraphPad Prism 10 software and are described in the figure legends. The p-values were designated as follows: *p<0.05; **p<0.01, ***p<0.001. Values of p > 0.05 were considered not significant (n.s.).

### 2.11 Global proteome analysis

For proteome profiling, HEL cell pellets were lysed in urea lysis buffer (8 M urea, 20 mM HEPES, pH 8.0, 1 mM sodium orthovanadate, 2.5 mM sodium pyrophosphate, 1 mM beta-glycerophosphate). Protein concentrations of the lysates were determined using the 660 nm assay kit (ThermoFisher Scientific) according to the manufacturer’s instructions. 10 µg of protein per sample were reduced with DTT (10 mM for 1 h at 37 °C), alkylated with iodoacetamide (25 mM for 15 min at 37 °C in the dark) and digested using Lys-C (Wako/Fujifilm) for 2 h at 37 °C in an enzyme-to-substrate ratio of 1:50 (w/w). After dilution with 20 mM HEPES (pH 8.0) to a concentration of 2 M urea, digestion was continued overnight with trypsin (Promega) at 37 °C and 1:50 (w/w) enzyme-to-substrate ratio. The peptide mixtures were acidified, purified using C18 spin tips (Havard) and dried by vacuum centrifugation. Afterward, the peptide samples were dissolved in 0.1 % formic acid and peptide concentrations were determined using a fluorometric peptide assay (ThermoFisher Scientific). The peptide samples were analyzed by LC-MS/MS on a Vanquish Neo UHPLC system (ThermoFisher Scientific) coupled online to an Orbitrap Astral mass spectrometer (ThermoFisher Scientific) in a data-independent acquisition scheme (DIA). 400 ng of peptides from each sample were concentrated and desalted on a PepMap Neo trap cartridge (ThermoFisher Scientific, particle size 100 Å, inner diameter 300 µm, length 5 mm), followed by separation on a 15 cm PepMapNeo C18 analytical column (ThermoFisher Scientific) using a 30 min method (27 min linear gradient) of 1% to 28% acetonitrile in 0.1% formic acid at a flow rate of 800 nl/min. Precursor ion survey scans were acquired using the Orbitrap mass analyzer with the following parameters: resolution 240,000, scan range *m/z* 380-980, automatic gain control (AGC) target 5x10^6^, maximum injection time 10 ms, RF lens setting 40%. For fragment ion scans using the Astral mass analyzer, precursor ions were isolated for collision-induced dissociation (HCD) through each survey scan with an isolation window of *m/z* 2, resulting in 299 scan events. The normalized HCD collision energy was set to 25% and for fragment ion analysis the AGC target was 5x10^4^ at a maximum injection time of 3 ms.

Raw DIA data were analyzed using Proteome Discoverer (version 3.1.1.93, ThermoFisher Scientific). Spectra were searched against the Uniprot human reference proteome and 245 frequently observed contaminants using the CHIMERYS search algorithm. The mass tolerance for fragment ions was set to 10 ppm. Oxidation of methionine was considered as dynamic modification while carbamidomethylation of cysteine was defined as a fixed modification. The peptide length was defined to be between seven to 30 amino acids with one allowed missed cleavage site. One to four charges per peptide were allowed. At both peptide and protein level, the false discovery rate (FDR) was set at 1%. Further data processing was done using R studio (version 2024.09.1). First, contaminants were removed. To correct for sample loading and technical variation, protein intensities were first scaled to the median of total sample intensities (sample loading normalization), followed by median normalization to align sample medians to the global median. Abundances were log_2_-transformed and proteins with >30% missing values across samples were excluded from analysis. The remaining missing values were imputed from a pre-defined normal distribution. Fold changes were calculated as the difference between the mean log_2_ intensities of each condition (n = 3 per group) and statistical significance was assessed using a two-sided t-test with Benjamini-Hochberg correction. (Supplementary table S2, data provided online (MassIVE Repository: Daten: https://doi.org/doi:10.25345/C5PV6BM4R (BSA_Glc)

### 2.12 Seahorse measurements

For Seahorse measurement the AML cell line HEL and primary AML blasts (0.5*10^6^ cells/mL) were cultured for 24 hours in RPMI 1640 medium w/o glucose (#11879020, Gibco/ThermoFisher) supplemented with 10% fetal calf serum (FCS, Sigma-Aldrich/Merck) and 100 IU/ml penicillin and 100 mg/ml streptomycin (P/S, #15140122, Gibco/ThermoFisher) and 0.5 g/L or 2 g/L glucose (D-glucose, #A2494001, Gibco/ThermoFisher) at 37°C in a humidified 5% CO_2_ incubator. For bovine serum albumin addition 5% (w/v) BSA (#A9418, Sigma-Aldrich/Merck) was added to the respective medium and the pH value was adjusted. For primary AML blasts cytokines were added in the following concentrations: 25 ng/ml hTPO (#300-18, Peprotech), 50 ng/ml hSCF (#300-07, Peprotech), 20 ng/ml hFLT3-Ligand (#300-19, Peprotech), 20 ng/ml hIL3 (#200-03, Peprotech).

Subsequently, the cellular ATP production was analyzed by ATP rate assays on a Seahorse XFe96 extracellular flux analyzer (Agilent). 50,000 cells were seeded on Seahorse 96-well cell culture plates at the day of measurement and equilibrated for 30 min before measurement in XF RPMI buffer (Agilent) supplemented with 0.5 mM or 2 mM L-glucose and 1 mM L-glutamine (all from Agilent). Cells were treated with 2.5 µM oligomycin and 0.5 µM rotenone and antimycin A (all from Cayman Chemicals) prior measurement of oxygen consumption rate and extracellular acidification.

### 2.13 CRISPR/Cas9, shRNA and production of lentiviral pseudotyped particles

Single guide RNAs (sgRNAs) targeting human GS were designed with a CRISPR design web-interface (https://benchling.com). The human GS knock-out (KO) vector was obtained by cloning the annealed target-specific oligonucleotides listed in table 3 into the BsmBI site of the pLentiCRISPRv2EGFP_ΔCas9 plasmid, a derivative of pLentiCRISPRv2 (Addgene, #52961) using the Golden Gate protocol [34]. A vector carrying a non-target control (NTC1) sequence was used as control (Table 2). After cloning, plasmids were purified and verified by sequencing. As shown in table 3, human and mouse MISSION^®^shRNA vectors targeting GLUL were obtained as bacterial stocks from Merck/Sigma-Aldrich/Merck. For control MISSION pLKO-1-puro non-target shRNA Control (Merck/Sigma-Aldrich/Merck, #SHC016) targeting no known genes from any species was used (shNTC).

**Table 2:**
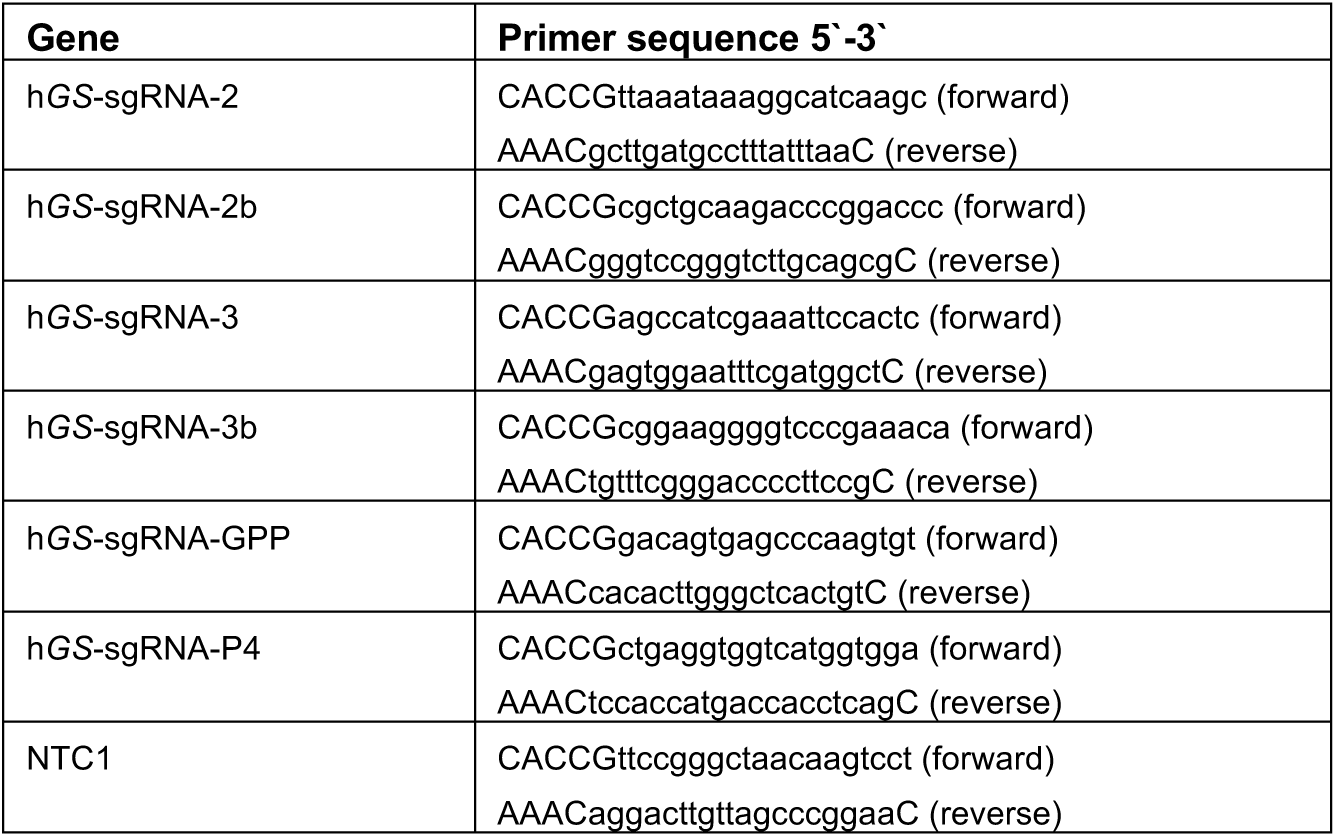
Oligonucleotides for sgRNAs.

**Table 3:**
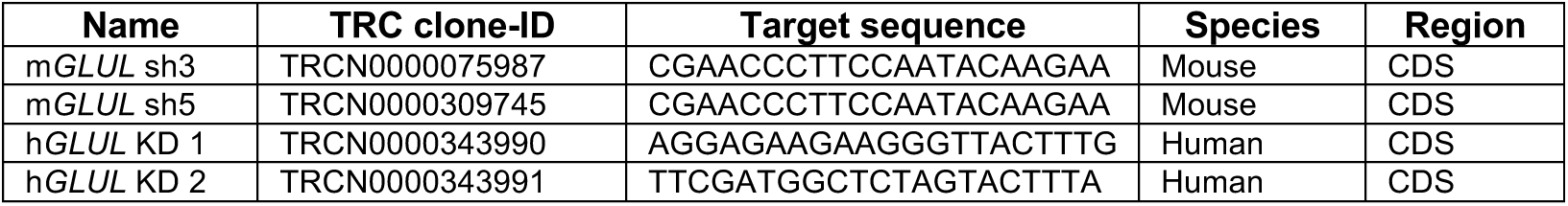
MISSION shRNAs.

Lenti-X 293T cells (Takara, #Z2180N) were used to produce viruses [35] using the packaging plasmids pMD2.G (Addgene, #12259) and psPAX2 (Addgene, #12260). Transduction efficiency was measured five days post transduction by flow cytometry using a BD LRS Fortessa device (Becton Dickinson). Data was analyzed with the BD FACSDiva software (Becton Dickinson).

### 2.14 Global Metabolomics using Ultrahigh Performance Liquid Chromatography-Tandem Mass Spectroscopy (UPLC-MS/MS) (Metabolon Inc.)

HEL cells were transduced with lentiviral particles carrying the control sgRNA (NTC1) and the sgRNA targeting GS (sgRNA GPP) (Table 2) and subdivided into five technical replicates. Four days post transduction, HEL NTC and GS KO cells were cultured in normal RPMI medium or with RPMI medium containing 5% BSA (see section 2.1) for 24 hours, harvested and washed with ice-cold 1xPBS. Cell pellets were frozen in liquid nitrogen and stored at −80°C until shipment. The same experiment was performed in RPMI medium containing reduced glucose concentration (0.5 g/L) (see also 2.1). Sample extraction, UPLC-MS/MS analysis, and initial data processing were performed by Metabolon Inc (617 Davis Drive, Suite 100, Morrisville, NC 27560, USA) (Supplementary table S3). Due to insufficient knockout efficiency, GS-KO samples were excluded from downstream analysis (data not shown). Batch-normalized metabolite data (provided by Metabolon) were further processed in Python 3.10.12 using the following libraries: *pandas* 2.2.2, *numpy* 1.26.4, *scipy* 1.13.0, *statsmodels* 0.14.1, *scikit-learn* 1.4.2, *matplotlib* 3.8.4, and *IPython* 8.24.0. Missing values were imputed using the row-wise median, followed by natural-log transformation. Metabolites for which more than two replicates per group required imputation were excluded to avoid artefacts (Supplementary table S3).

Principal component analysis (PCA) was performed on z-scored metabolite intensities to assess global metabolic variation. Differential abundance analysis comparing NTC_SS vs NTC_BSA was performed using Welch’s t-tests, with multiple-testing correction by the Benjamini–Hochberg procedure. Fold changes were calculated from group-wise medians on the linear scale. Analysis scripts are available from the corresponding author upon reasonable request.

Pathway enrichment analysis was carried out in MetaboAnalyst v6.0 using significantly regulated metabolites (p < 0.05) after quality control. Separate KEGG- and SMPDB-based analyses were performed. Bubble plots were generated with –log₁₀(p-value) (y-axis) and pathway impact (x-axis); bubble color denotes significance and bubble size represents enrichment ratio. Where necessary, manual corrections were made (e.g., substitution of D-amino acid entries with their corresponding L-forms), while retaining the original pathway statistics from MetaboAnalyst (Supplementary Table S3).

Metabolite annotation was based primarily on the Human Metabolome Database (HMDB v5.0). For compounds lacking unique HMDB IDs, a custom mapping strategy was applied using parent HMDB compound classes to enable pathway-level assignment.

### 2.15 Ammonia measurement in cell culture supernatants

To measure the ammonia concentration in cell culture supernatants upon BSA feeding, cells (0.5*10^6^ cells/mL) were cultured for 24 hours in RPMI 1640 medium w/o glucose (#11879020, Gibco/ThermoFisher) supplemented with 10% fetal calf serum (FCS, Sigma-Aldrich/Merck) and 100 IU/ml penicillin and 100 mg/ml streptomycin (P/S, #15140122, Gibco/ThermoFisher) and 0.5 g/L or 2 g/L glucose (D-glucose, #A2494001, Gibco/ThermoFisher) at 37°C in a humidified 5% CO_2_ incubator. For bovine serum albumin addition 5% (w/v) BSA (#A9418, Sigma-Aldrich/Merck) was added to the respective medium and the pH value was adjusted. For primary AML blasts cytokines were added in the following concentrations: 25 ng/ml hTPO (#300-18, Peprotech), 50 ng/ml hSCF (#300-07, Peprotech), 20 ng/ml hFLT3-Ligand (#300-19, Peprotech), 20 ng/ml hIL3 (#200-03, Peprotech). Supernatant samples were measured with an Ammonia assay kit (#ab83360, Abcam) according to the manufacturer’s instructions.

### 2.16 Ammonium, albumin and glucose measurements in patient samples

In order to measure ammonium levels in patients with AML, peripheral blood samples and bone marrow aspirates were taken from patients at the time of initial diagnosis and during the treatment course when remission status was controlled. As negative controls (bone marrow without malignant infiltration), samples from lymphoma patients were used. These samples were obtained during routine diagnostic bone marrow aspiration and showed no evidence of infiltration upon cytological, flow cytometric, and histological evaluation. As an internal reference we used glucose levels in peripheral blood and bone marrow aspirate. During the procedure, 8-10 mL of bone marrow blood were aspirated in a syringe (#4606108V, B.Braun, Germany) without any anticoagulant. The aspirate was immediately transferred to an EDTA monovette (ammonium) or to a serum monovette (glucose, albumin) (EDTA: 05.1167; Serum: 01.1602, Sarstedt AG, Germany), respectively. The EDTA monovette was transported at 2–8 °C, and both monovettes were sent immediately after sample collection to the Central Laboratory of Goethe University Medicine in Frankfurt, Germany. Albumin, glucose and ammonium levels were measured on the Cobas® 8000 automated analyzer (Roche Diagnostics, Basel, Switzerland) as part of their routine diagnostics. Patient data can be found in Supplementary table S1 (patient data used in ammonia cohort).

### 2.17 Targeted metabolomics using Gas Chromatography-Mass Spectroscopy (GC-MS): Intracellular tracking of ^15^NH_4_Cl

A total of 10x10^6^ HEL cells (1x10^6^ cells/mL) were cultured in RPMI 1640 medium w/o glutamine (#21870076, Gibco/ThermoFisher) supplemented with 10% dialyzed fetal bovine serum (dFBS, #26400044, Gibco/ThermoFisher) and 100 IU/ml penicillin and 100 mg/ml streptomycin (P/S, #15140122, Gibco/ThermoFisher). For Gln-replete condition 300mg/L L-Gln (L-glutamine 200mM, #A2916801, Gibco/ThermoFisher) was added. After one hour, cells were treated with 5 mM ^15^NH_4_Cl (#299251, Sigma-Aldrich/Merck) for 1, 6 and 24 hours. In Gln-replete condition, only the 24 hours sample was prepared. At the end of the experiment, cells were counted, media and cell pellets were separated. Cells were washed twice with ice-cold PBS before snap-freezing with liquid nitrogen and were stored at −80°C until sample processing. We extracted polar intracellular metabolites from pellets (10x10^6^ cells/sample) using ice-cold MeOH:H_2_O:CHCl_3_ (1:1:2) extraction method in 16 mm diameter glass tubes. 5 ng of norvaline was added to each sample as recovery standard for relative quantification of metabolites. Tubes were shaken for 30 minutes at 4°C and then were centrifuged for 15 minutes at 3100 rcf at 4°C. The upper polar phase containing the polar metabolites was transferred to another tube and was dried under air flow. 50 μl of DCM (dichloromethane) were added to each tube and dried with nitrogen gas to guarantee total dehydration of the sample prior to derivatization. The interphase containing proteins was also transferred to another tube. The interphase was washed with cold MeOH, then dried under air flow and next disaggregated using 30% KOH solution at 100°C for 15 minutes. Protein content was measured using BCA assay kit (#23227, Pierce/ThermoFisher).

Derivatization of samples containing polar metabolites was run as follows: samples were firstly derivatized using 2% methoxamine hydrochloride in pyridine for 90 minutes with heating at 37°C and shaking. As a follow up, samples were incubated for 1 h at 55 °C with MBTSTFA+ 1% TBDMCS, and were then transferred to Gas Chromatography-Mass Spectrometry (GC-MS) vials. GC-MS analysis was run under electron impact ionization using a 7890A GC equipped with a HP5 capillary column coupled to an Agilent 5975C MS (Agilent, Santa Clara, CA, USA). For all GC-MS measurements, 2 μL of sample was injected at 270°C using pulsed split mode. GC oven temperature was held at 100°C for 3 min after injection and increased to 300°C through four consecutive ramp temperatures: first to 165°C at 10°C/min, next to 225°C at 2.5°C/min, next to 265°C at 25°C/min and finally to 300°C at 7.5°C/min. All spectra were recorded using full scan MS configuration, from m/z 100 to 650. Supplementary table S4 shows the retention time (RT) and m/z values that were monitored in our analysis. All integrations were performed manually, considering all the peak area for each metabolite. Relative concentration of metabolites in cell pellets (Metabolite ion counts) were normalized to norvaline ion counts as recovery standard and to the protein content. Midcor software was used to correct natural enrichment and ion interferences in the measured isotopologue distributions and to quantify ^15^N labelling incorporation and real isotopologue distributions [36]. M+1 and M+2 indicate how many ^15^N were incorporated in the analyzed amino acids.

### 2.18 ^1^H-^15^N heteronuclear NMR experiment

HEL cells were transduced with lentiviral particles carrying the control sgRNA (NTC1) and the sgRNA targeting GS (sgRNA GPP) (Table 3) and subdivided into two technical replicates. Four days post transduction HEL NTC and GS KO cells were non-treated or treated with 2% (w/v) of ^15^N-labelled Arthrospira maxima hydrolysate (Silantes, Germany) in RPMI medium supplemented with 1% FCS for 24 hours. Cells were spun at 2000 rpm for 4 minutes at 4°C, the supernatant was discarded, cells were re-suspended in 1 ml ice cold 1xPBS and centrifuged for 30 seconds at 10000 rcf at 4°C. The supernatant was discarded and the pellet was resuspended in 400 μl MeOH (HPLC grade) pre-chilled on dry ice. Samples were then transferred to glass vials and 325 μL ddH_2_O and 400 μl chloroform (HPLC grade), pre-chilled on wet ice, was added. Samples were vortexed for 30 seconds prior to resting for 10 minutes and subsequent centrifugation for 15 minutes at 4000 rpm. 400 µL from the polar phase was removed using a glass Hamilton syringe and dried using a SpeedVac concentrator and stored at −80°C until further analysis.

The dried metabolite extracts were resuspended in 60 µL of NMR buffer (100 mM sodium phosphate buffer containing 0.5 mM TMSP (Na^+^ salt of trimethylsilylproprionat) in 10% D_2_O at pH 7.0) and sonicated for 10 minutes. For media samples, 10% D_2_O was added to each sample directly. Samples were then transferred to *Shigemi* 3 mm NMR tubes (Cortecnet, France) and measured on an AV950 MHz Bruker spectrometer equipped with a 5 mm TCI cryoprobe (Bruker, Switzerland). 2D ^1^H-^15^N heteronuclear single quantum coherence (HSQC) experiments were performed at 25°C using a Bruker library pulse program (hsqcetf3gp) with 25% non-uniform sampling acquisition mode (25% NUS), 1024 number of scans, a TD of (2048, 110) and a spectral width of 8279 Hz. Spectral resolution was 150 Hz for the indirect dimension. Data analysis was performed in TopSpin 3.5 (Bruker, Germany).

### 2.19 Proteome and transcriptome meta-analysis

For integration of primary patient transcriptome data of enzymes central to cellular ammonia detoxification pathways, the datasets from Bloodspot (servers.binf.ku.dk/bloodspot/) MILE study (GSE13159) [37, 38] and human normal hematopoiesis (GSE42519) were used. The suitability of the probe sets used were further analyzed using the ENSEMBL genome browser (ensemble.org). Only probe sets that were completely positioned within exonic regions of the respective genes were considered for the subsequent analyses (Table 4). Fold changes (f.c.) were calculated according to log2-transformed expression values compared with normal HSPCs.

**Table 4:** Probe set analysis MILE study (GSE13159).

| Gene | Probe set | Targeting region | Meta-analysis |
| --- | --- | --- | --- |
| GLUD1 | 200946_x_at | 3' UTR | yes |
|  | 200947_s_at | 3' UTR | yes |
| GLUL | 215001_s_at | Far in 3' UTR | No |
|  | 217202_s_at | Exons 4, 6 | yes |
|  | 200648_s_at | Exon 7, 3' UTR | yes |
|  | 242281_at | 3' UTR of ENST00000331872, not targeting ENST00000339526 | No |
| CPS1 | 204920_at | 3' UTR | yes |
|  | 217564_s_at | 3' UTR | yes |

For the integration of proteomic data of the respective ammonia detoxifying enzymes, datasets from De Boer et al. 2018 [39] (PRIDE: PXD030463), Kramer et al. 2022 [40] (MassIVE: MSV00089012) and our own published data set from Jayavelu et al 2022 [41] (PRIDE: PXD023201 and PXD028007, Supplementary table S5) were analyzed. Fold changes (f.c.) were calculated according to log_2_-transformed or log10-transformed expression values compared with normal mononuclear cells (MNC) or CD34-positive cells.

*GLUL* gene expression in 1450 cell lines derived from 74 primary diseases was analyzed using the public database DepMap portal (depmap.org) provided by the Broad Institute (Supplementary table S5). Gene expression TPM values are reported after log2 transformation

### 2.20 Colony formation assay

Colony-forming cells were assayed in methylcellulose (MethoCult H4100, #04100 for THP-1 cells and MethoCult M3534, #03534 for mouse myeloid progenitor/MN1 cells; StemCell Technologies). For mouse myeloid progenitors, methylcellulose was supplemented with 10 ng/mL mIL-3 (#213-13, Peprotech), 10 ng/mL hIL-6 (#200-06, Peprotech), 50 ng/mL mSCF (#250-03, Peprotech). For each plating 500 viable cells/ tissue culture dish (TC dish, standard, with Grid, #83.3900.002, Sarstedt) were plated in triplicates and incubated at 37°C under 5% (v/v) CO_2_. Colonies were evaluated microscopically 7 days (THP1) or 4 days primary myeloid progenitor MN1) after plating by using standard criteria. For replating, cells were eluted from the methylcellulose, washed, counted and replated as described above.

To evaluate the leukemic potential of primary AML bone marrow mononuclear cells (MNC), leukapheresis samples as well as sorted CD45^dim^/CD34^+^/CD38^-^ leukemia initiating cells (LICs), cells were assayed in methylcellulose (MethoCult H4435 Enriched, #04435, StemCell Technologies) supplemented with 25 ng/ml hTPO (#300-18, Peprotech), 50 ng/ml hSCF (#300-07, Peprotech), 20 ng/ml hFLT3-Ligand (#300-19, Peprotech), 20 ng/ml hIL3 (#200-03, Peprotech). For each plating, 5000/10000 viable cells (MNC) or 1000 LICs/ tissue culture dish (TC dish, Standard, with Grid, #83.3900.002, Sarstedt) were plated in duplicates and cultured in a hypoxia chamber (Biospherix, X3 Xvivo System, USA). Cells were kept at 37°C under 1% O_2_, 5% CO_2_ and 94% N_2_. Colony forming units (CFU-L) were evaluated microscopically 21 days after plating by using standard criteria. To analyze the effects of GS inhibition methylcellulose was supplemented with 0.5, 1 and 2.5 mM MSO. For studying the molecular function of GS in primary AML cells, methylcellulose was supplemented with 2mM NH_4_Cl (Ammonium chloride, #A9434, Sigma-Aldrich/Merck).

### 2.21 Xenograft mouse model

Ten-to fourteen-week-old, non-irradiated NSG (NOD.Cg-Prkdcscid Il2rgtmWjl/SzJ) mice were transplanted intravenously with 3*10^6^ THP-1 cells transduced with either sh*NTC* or h*GLUL* KD 1/2 shRNAs (see 2.13) four days prior transplantation. Mice were followed for overall survival analysis and sacrificed when mice showed severe signs of illness. Survival analysis was performed by the Kaplan-Meier method using GraphPad Prism (v9.5.1, GraphPad Software, Inc., San Diego, CA, USA) and the p-value was obtained by GehanBreslow-Wilcoxon test. Mouse organs including BM, spleen and liver as well as peripheral blood were isolated and analyzed for human CD33 and CD45 protein expression by flow cytometry (see 2.2., Table 1). The study was approved by the local German government (Regierungspraesidium Darmstadt) in Hesse, Germany (FK/1117).

### 2.22 Murine MN1-driven leukemia model

Bone marrow (BM) cells were isolated from C57BL/6J wild-type mice followed lineage depletion was performed using a mouse Lineage Cell Depletion Kit (#130-110-470, Miltenyi Biotec). Transformation with pSF91-*MN1*-IRES-eGFP was performed as described previously [42]. Transformed MN-1 cells were cultured in (DMEM 4.5 g/L D-glucose, Gibco/ThermoFisher, #41965-039) supplemented with 15% fetal calf serum (FCS, Sigma-Aldrich/Merck), 100 IU/ml penicillin and 100 mg/ml streptomycin (P/S, Gibco/ThermoFisher, #15140122), 10 ng/ml murine interleukin IL-3 (#213-13, Peptrotech), 10 ng/ml human IL-6 (#200-06, Peprotech) and 100 ng/ml murine SCF (#250-03, Peprotech) at 37°C in a humidified 5% CO_2_ incubator. Upon transformation MN-1 cells were transduced with either sh*NTC* or m*GLUL* sh3/sh5 shRNAs (see 2.7) and selected using puromycin for two days. Two hours before intravenously transplantation of 2*10^5^ transduced MN-1 cells, mice were non-lethally irradiated with 5.5 Gy. Mice were followed for overall survival analysis and sacrificed when mice showed severe signs of illness. Survival analysis was performed by the Kaplan-Meier method using GraphPad Prism (v9.5.1, GraphPad Software, Inc., San Diego, CA, USA) and the p-value was obtained by GehanBreslow-Wilcoxon test. Mouse organs including BM and spleen as well as peripheral blood were isolated and GFP-positive cells were analyzed for lineage positive/negative cells and murine CD117, murine CD11b, murine Ly-6G, murine Ly6A/E, murine CD16/32 and murine CD34 protein expression by flow cytometry (see 2.2., Table 1). The study was approved by the local German government (Regierungspraesidium Darmstadt) in Hesse, Germany (FK/1117).

### 2.23 Free amino acid measurements in peripheral blood and bone marrow of AML patients

The concentration of free amino acids was measured in bone marrow (BM) samples and peripheral blood (PB) of AML patients at initial diagnosis. PB was drawn on the same day during routine testing into an EDTA monovette. BM blood was obtained through aspiration into a syringe without anticoagulants (#4606108V, B.Braun, Germany) and immediately transferred to an EDTA monovette (EDTA: 05.1167, Sarstedt AG, Germany). In order to prevent extracorporal hemolysis, samples were centrifuged within 20 minutes after aspiration for at least 10 minutes at 400 x g. The supernatant was transferred to either Eppendorf tubes and stored at −80°C until further handling or transferred to screw cap tubes (62.612, Saarstedt AG, Nümbrecht) for immediate measurement in the laboratory of the Hessische Kindervorsorgezentrum, University Medicine Frankfurt, Germany.

Amino acid analysis was performed using a Biochrom 30+ amino acid analyzer equipped with a two-wavelength photometer and flow cell (440/570 nm) and an Alias autosampler (Biochrom, UK). Data acquisition and evaluation were carried out using Agilent OpenLab EZChrom software (Version A.04.10, Build 04.10.32).

## 3. Results

### AML cells acquire extracellular protein through macropinocytosis and lysosomal proteocatabolism

It is well established that several amino acids (Met, Cys, Gln, and branched-chain amino acids) are critical for AML cell survival and proliferation [10, 11, 13, 16, 17]. To investigate how AML cells meet their anabolic demands given this dependency, we explored whether they engage in extracellular protein scavenging as an alternative nutrient acquisition strategy within the BM microenvironment.

We measured the concentration of free amino acids in BM plasma from eleven diagnosed AML patients at initial diagnosis. Most amino acids, with the exception of asparagine (Asn, N) and histidine (His, H), were within the expected physiological range for blood plasma (Fig. 1A, left). However, paired analysis of BM versus peripheral blood (PB) plasma revealed that levels of multiple amino acids, including alanine (Ala, A), isoleucine (Ile, I), leucine (Leu, L), proline (Pro, P), valine (Val, V), phenylalanine (Phe, F), tryptophan (Trp, W), tyrosine (Tyr, Y), lysine (Lys, K), serine (Ser, S), threonine (Thr, T), and Gln, were significantly lower in the BM (Fig. S1). These findings suggest that, despite being physiologically nourished, the BM microenvironment imposes localized amino acid constraints that may require adaptive metabolic strategies by AML cells.

**Figure 1.**
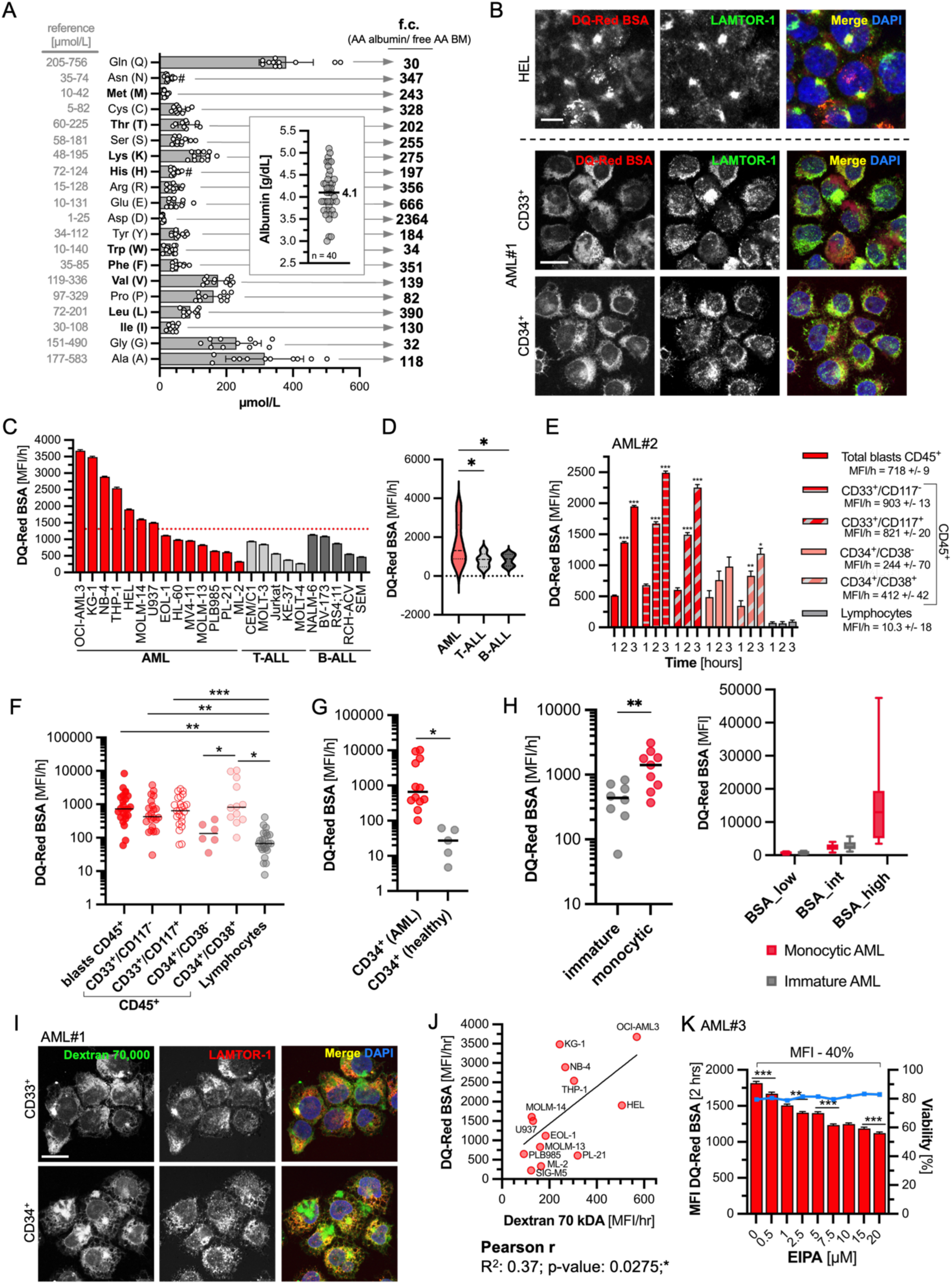
AML cells acquire extracellular protein through macropinocytosis and lysosomal proteocatabolism. (A) Calculation of free amino acids derived from the complete conversion of serum albumin at a concentration of 4.1 g/dL compared to the concentration of free amino acids in the bone marrow (BM) of AML patients at first diagnosis (fold change). Bar graph: free amino acid measurements in the BM of nine AML patients at diagnosis, shown in µmol/L. Left: serum reference free amino acid concentrations. Dot plot: albumin concentrations measured in peripheral blood (PB) from 40 AML patients at diagnosis, shown in g/dL. (B) Confocal images of HEL cells (upper panel) and primary AML CD33⁺ and CD34⁺ blasts treated for 3 and 5 hours, respectively, with a final concentration of 10 µg/mL DQ-Red bovine serum albumin (DQ-Red BSA). Lysosomes were stained using a primary antibody directed against LAMTOR1. Nuclei were stained with DAPI. Scale bar: 10 µm. (C) Protein catabolism rate measured by mean fluorescence intensity per hour (MFI/h) of DQ-Red BSA in fourteen AML (red bars), five T-cell acute lymphoblastic leukemia (T-ALL, light grey bars), and five B-cell acute lymphoblastic leukemia (B-ALL, dark grey bars) cell lines. Bar graph represents MFI/h ± SEM (left). Red dotted line indicates the median of the MFI/h of the AML cell lines. (D) Violin plots comparing protein catabolism rates (MFI/h) in AML, T-ALL, and B-ALL cell lines. Brown–Forsythe and Welch ANOVA with Dunnett’s post hoc test. **, p < 0.01. (E) MFI of DQ-Red BSA in primary AML cell populations after 1, 2, and 3 hours. Bar graphs represent mean ± SEM. Two-way ANOVA with Bonferroni’s post hoc test. *, p < 0.05; **, p < 0.01; **, p < 0.001. (F) Protein catabolism rate (MFI/h of DQ-Red BSA) in primary AML cell populations and matched lymphocytes from 24 AML patients at first diagnosis. Each cell population was compared to lymphocytes using an unpaired t-test with Welch’s correction (, p < 0.05; **, p < 0.01; *, p < 0.001). (G) Protein catabolism rate (MFI/h of DQ-Red BSA) in primary CD34⁺ AML blasts and CD34⁺ cells derived from healthy donors. Unpaired t-test with Welch’s correction (, p < 0.05). (H) (Left) Protein catabolism rate (MFI/h of DQ-Red BSA) in immature blasts compared to blasts with monocytic differentiation. Unpaired t-test with Welch’s correction (, p < 0.05). (Right) Mean DQ-Red BSA fluorescence intensity (MFI) of flow cytometrically defined DQ-BSA_low, DQ-BSA_int, and DQ-BSA_high subpopulations across AML samples classified as immature (grey) or monocytoid-differentiated (red). Immature AML samples showed no DQ-BSA_high subpopulation. (I) Confocal images of primary AML CD33⁺ and CD34⁺ blasts treated for 6 hours with a final concentration of 0.05 mg/mL dextran (Oregon Green 488, 70,000 MW). Lysosomes were stained using a primary antibody directed against LAMTOR1. Nuclei were stained with DAPI. Scale bar: 10 µm. (J) Pearson correlation between protein catabolism rate (DQ-Red BSA [MFI/h]) and macropinocytosis rate (dextran 70,000 [MFI/h]) across 13 AML cell lines. (K) MFI of DQ-Red BSA (red bars) and cell viability (%) in primary AML blasts pretreated for 24 hours with increasing concentrations of EIPA. Statistics for MFI of DQ-Red BSA: one-way ANOVA with Bonferroni’s post hoc test. **, p < 0.01; ***, p < 0.001.

To assess the relevance of extracellular protein as a nutrient reservoir, we next quantified albumin concentrations in PB plasma from 40 AML patients at diagnosis. The median albumin concentration was 4.1 g/dL (Fig. 1A, dot plot). Using this value as a basis, we calculated the theoretical amino acid yield resulting from complete proteolytic degradation of albumin. Based on albumin’s amino acid composition, full degradation would generate amino acid amounts corresponding to concentrations approximately 30-to 2,345-fold higher than those of free amino acids measured in serum (Fig. 1A, middle and right).

To test whether AML cells actively catabolize extracellular protein, we used dye-quenched (DQ) Red-labeled bovine serum albumin (DQ-Red BSA) as a reporter substrate, given that albumin is the most abundant protein in human plasma, comprising approximately 60% of total plasma protein [43]. Confocal microscopy of the AML cell line HEL, as well as sorted CD33⁺ and CD34⁺ primary AML blasts, revealed strong intracellular fluorescence co-localizing with LAMTOR1, a lysosomal marker indicative of active uptake and degradation of extracellular albumin (Fig. 1B). Quantification of mean fluorescence intensity per hour (MFI/h) across multiple AML cell lines confirmed consistently higher rates of DQ-Red BSA processing compared to T-cell and B-cell acute lymphoblastic leukemia (T-ALL and B-ALL) cell lines (Fig. 1C/D; Fig. S2). In primary AML samples, DQ-Red BSA degradation was significantly higher in distinct leukemic blast populations than in matched lymphocytes (Fig. 1E/F), and also exceeded that observed in healthy hematopoietic progenitor (CD34⁺) cells (Fig. 1G).

Interestingly, flow cytometric analysis of DQ-Red BSA uptake in primary AML samples revealed that more differentiated blast subpopulations (associated with higher side scatter (SSC) and CD45 expression) exhibited significantly higher DQ-Red BSA fluorescence intensity than immature blast populations. By integrating these data with routine immunophenotyping, we found that AML cases with immunophenotypic features of monocytic differentiation consistently showed enhanced extracellular protein catabolism compared to those with immature phenotypes (Fig. 1H, Fig. S3). These findings suggest that protein scavenging is particularly prominent in AML subtypes with monocytic features, potentially reflecting lineage-specific metabolic rewiring.

In pancreatic cancer, enhanced extracellular protein uptake has been attributed to macropinocytosis, a process driven by oncogenic KRAS mutations [21, 22, 44]. Although KRAS alterations are uncommon in AML (approximately 5-12% of AML cases [1, 45], we hypothesized that AML cells may possess intrinsic macropinocytic activity due to their hematopoietic origin. To evaluate this, we exposed sorted CD33⁺ and CD34⁺ primary AML blasts to fluorescently labeled 70 kDa dextran, a classical substrate for measuring macropinocytosis, and counterstained with LAMTOR1 as a lysosomal marker. Confocal imaging revealed robust dextran uptake with clear lysosomal co-localization, consistent with macropinocytic internalization and processing. To further determine whether macropinocytosis contributes to extracellular protein scavenging, we quantified dextran uptake and DQ-Red BSA processing across 13 AML cell lines. This analysis revealed a significant positive correlation between the two (Fig. 1J), suggesting that protein catabolism in AML is mechanistically linked to macropinocytic activity. Details of the dextran uptake analysis are provided in Fig. S4A.

To functionally confirm the role of macropinocytosis in AML protein scavenging, we made use of 5-(N-ethyl-N-isopropyl) amiloride (EIPA), a Na^+^/H^+^ exchanger (NHE) inhibitor that blocks macropinocytosis. EIPA was applied across a concentration range that did not impair cell viability when cells were pre-treated for 24 hours prior to DQ-Red BSA exposure. As shown in figure 1K, in primary AML blasts, EIPA treatment led to a dose-dependent reduction in DQ-Red BSA MFI by 40%, indicating diminished extracellular protein catabolism (Fig. 1K). Similar results were observed in the AML cell line HEL, where EIPA reduced DQ-Red BSA processing without affecting viability (Fig. S4B/C). Together, these findings support macropinocytosis as an important mechanism contributing to extracellular protein catabolism in AML.

### Extracellular protein catabolism sustains energy metabolism and engages nitrogen-handling pathways in AML

To assess whether AML cells can metabolically exploit extracellular protein to support cell growth, we supplemented cultures with 5% BSA. Across multiple AML cell lines (HEL, THP-1, MOLM-13, and MV4-11), BSA significantly enhanced proliferation (Fig. 2A). This proliferative benefit was preserved in human plasma-like medium (HPLM), which mimics physiological concentrations of free amino acids (Fig. 2B). These findings suggest that protein catabolism supports leukemic cell growth under both nutrient-replete and physiologic amino acid conditions.

**Figure 2:**
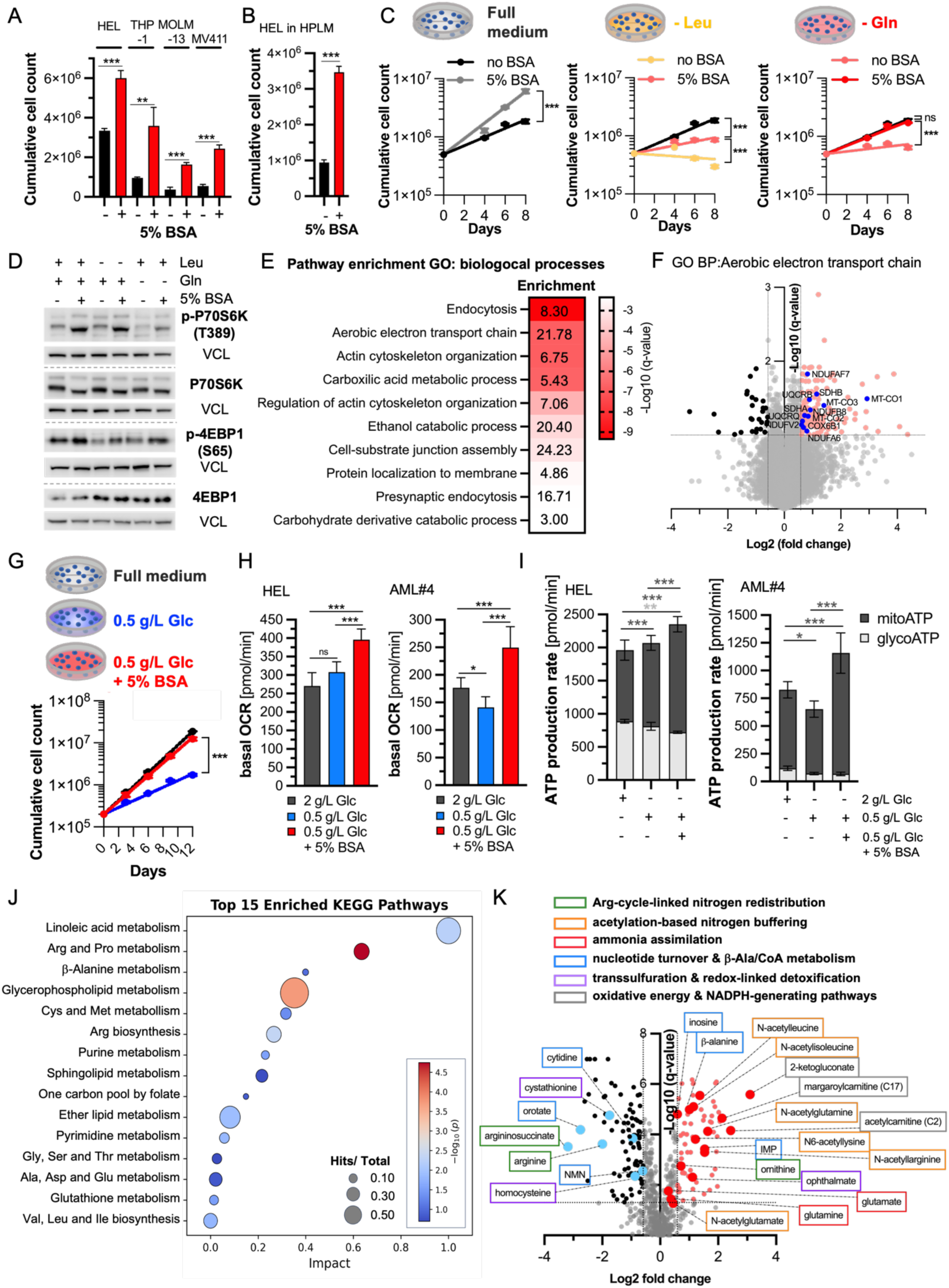
Extracellular protein catabolism sustains energy metabolism and engages nitrogen-handling pathways in AML. (A) Cumulative growth of HEL, THP-1, MOLM-13, and MV4-11 cells cultured in RPMI medium supplemented with 1% FCS, with or without 5% bovine serum albumin (BSA, w/v), over 6 days in triplicates. Unpaired t-test; **, p < 0.01; ***, p < 0.001. (B) Cumulative growth of HEL cells cultured in human plasma-like medium (HPLM) supplemented with 1% FCS, with or without 5% BSA (w/v), over 4 days in triplicates. Unpaired t-test; ***, p < 0.001. (C) Cumulative growth assay of HEL cells cultured in full medium, leucine-free (−Leu), or glutamine-free (−Gln) medium, with or without 5% BSA (w/v), for 8 days in duplicates. Extra sum-of-squares F test to evaluate differences in exponential growth-curve fitting; ***, p < 0.001. (D) Western blot analysis of HEL cells cultured in full, leucine-depleted (Leu), or glutamine-depleted (Gln) medium, with or without 5% BSA (w/v), for 24 hours. Detection of endogenous phospho-P70S6K (Thr389), total P70S6K, phospho-4EBP1 (Ser65), and total 4EBP1. Vinculin (VCL) served as loading control. (E) Pathway enrichment analysis (GO biological processes) of the global proteomics dataset from HEL cells cultured for 7 days in RPMI medium supplemented with 5% BSA (w/v). Numbers indicate enrichment score. Red color intensity represents −log₁₀(q-value). (F) Volcano plot highlighting enrichment (log₂ fold change) and significance (−log₁₀ q-value) of proteins belonging to the GO biological process “aerobic electron transport chain.” (G) Cumulative growth assay of HEL cells cultured in full medium containing 2 g/L glucose (Glc), 0.5 g/L Glc, or 0.5 g/L Glc supplemented with 5% BSA (w/v) in dupkicates. Extra sum-of-squares F test to evaluate differences in exponential growth-curve fitting; ***, p < 0.001. (H) Basal oxygen consumption rate (OCR, pmol/min) in HEL cells and primary AML blasts (AML#4) cultured for 24 hours in the presence of standard glucose (2 g/L Glc) or reduced glucose (0.5 g/L Glc), with or without 5% BSA (w/v). One-way ANOVA with Bonferroni’s post hoc test; ***, p < 0.001. (I) ATP production rate (pmol/min) in HEL cells and primary AML blasts (AML#4) cultured for 24 hours in the presence of standard glucose (2 g/L Glc) or reduced glucose (0.5 g/L Glc), with or without 5% BSA (w/v). Dark grey bars indicate mitochondrial-derived ATP (mitoATP); light grey bars indicate glycolysis-derived ATP (glycoATP). Two-way ANOVA with Bonferroni’s post hoc test; *, p < 0.05; **, p < 0.01; ***, p < 0.001. (J) Pathway enrichment analysis of the global metabolomics dataset from HEL cells cultured for 24 hours with 5% BSA (w/v), showing the top 15 enriched KEGG pathways. Bubble size represents the metabolite hit/total ratio. Red–blue color scale represents −log₁₀(p-value). (K) Volcano plot of significantly increased and decreased metabolites upon BSA treatment (log₂ fold change). Red and blue dots highlight representative metabolites from five cellular nitrogen-handling nodes (arginine-cycle–linked nitrogen redistribution; acetylation-based nitrogen buffering; ammonia assimilation; nucleotide turnover and β-alanine/CoA metabolism; transsulfuration- and redox-linked detoxification), as well as oxidative energy- and NADPH-generating pathways.

To determine whether albumin could compensate for specific nutrient limitations, we tested its ability to rescue cell proliferation in media lacking Gln or Leu, which are both essential for AML proliferation [10,16]. BSA supplementation fully restored proliferation in Gln-depleted medium and partially rescued growth under Leu depletion (Fig. 2C), indicating that the proteolytic breakdown of protein in the form of albumin provides a usable amino acid source.

Given the established role of exogenously supplied free amino acids as well as amino acids generated through lysosomal protein degradation, particularly leucine, in activating mechanistic target of rapamycin complex 1 (mTORC1) [46–48], we next examined whether BSA-derived amino acids influence mTORC1 activity in AML. Western blot analysis showed that BSA increased phosphorylation of two canonical mTORC1 targets: ribosomal protein S6 kinase beta-1 (p70S6K, Thr389) and eukaryotic translation initiation factor 4E-binding protein 1 (4EBP1, Ser65), under full amino acids, Leu-depleted, and Gln-depleted conditions (Fig. 2D). Quantification confirmed consistent upregulation of mTORC1 activity across conditions (Fig. S5A). Consistent with anabolic pathway activation, BSA also enhanced nuclear sterol regulatory element binding transcription factor 2 (SREBP2) accumulation and expression of its target squalene epoxidase (SQLE), suggesting increased sterol biosynthesis (Fig. S5B). Conversely, BSA reduced phosphorylation of transcription factor EB (TFEB) at Ser211 and Ser122 in both endogenous and overexpression systems, pointing to a potential derepression of lysosomal gene expression (Fig. S5C/D). Together, these findings demonstrate that extracellular protein catabolism activates mTORC1 signaling and engages multiple downstream anabolic programs.

To explore the broader consequence of extracellular protein catabolism on cellular mechanisms and energy metabolism, we performed a global proteome analysis after BSA exposure. Among proteins significantly upregulated (≥1.5-fold, q < 0.1), gene ontology enrichment revealed strong representation of pathways related to the biological processes of endocytosis, aerobic electron transport chain, and actin cytoskeleton organization (Fig. 2E). In line with the enrichment of oxidative phosphorylation, multiple subunits across respiratory complexes I–IV were upregulated, including NDUFAF7, NDUFB8, NDUFA6, and NDUFV2 (complex I), SDHB (complex II), UQCRB and UQCRQ (complex III), and MT-CO1, MT-CO2, MT-CO3, and COX6B1 (complex IV) (Fig. 2F). Similarly, a corresponding protein interaction network (Fig. S5E) identified clusters related to endocytosis, mitochondrial function, and actin dynamics. Together, these findings indicate that protein/BSA feeding elicits a coordinated proteomic program that enhances both nutrient uptake capacity and mitochondrial energy machinery, that suggests a functional coupling between extracellular protein catabolism and oxidative metabolism.

To directly test this link, we next assessed whether protein/BSA-derived substrates can support energy production. The proteolytic breakdown of proteins yields amino acids that not only serve as anabolic precursors but also provide carbon skeletons capable of fueling the tricarboxylic acid (TCA) cycle and sustaining ATP synthesis. In glucose-restricted medium (0.5 g/L), BSA supplementation fully restored cell proliferation (Fig. 2G), indicating that AML cells can metabolize protein-derived carbon backbones to maintain growth. Consistent with this, ATP-rate measurements showed increased mitochondrial oxygen consumption rate (OCR) in HEL cells and primary AML blasts upon BSA supplementation (Fig. 2H). While HEL cells maintained basal OCR across glucose conditions, primary AML blasts showed a glucose-dependent reduction in OCR, which was reversed by BSA. Seahorse-based ATP production analysis further confirmed that BSA enhances mitochondrial ATP (mitoATP) production in both cell types (Fig. 2I), underscoring the contribution of protein catabolism to oxidative energy metabolism.

To define how extracellular protein scavenging reshapes global metabolism, we conducted untargeted metabolomics (UPLC-MS/MS) of HEL cells with and without BSA supplementation. The dataset comprises a total of 819 biochemicals, 781 compounds of known identity and 38 compounds of unknown structural identity. After quality filtering to exclude features with excessive imputation, 722 metabolites were retained for quantitative analysis. Principal component analysis (PCA) of the metabolomic profiles (Fig. S5F) revealed a clear separation between untreated control (Control) and albumin-supplemented (+BSA) samples. The first two principal components explained a cumulative 63.5% of total variance (PC1 = 48.9%, PC2 = 14.6%). Variation along PC1 primarily distinguished BSA-treated from control samples, indicating a coordinated and robust metabolic response to extracellular protein availability. The smaller contribution of PC2 reflects intra-group variability, which remained low relative to the treatment effect. Differential metabolite abundance analyses (Welch’s *t*-test with Benjamini–Hochberg FDR correction) revealed significant regulation of amino acid, lipid, nucleotide, and nitrogen-handling metabolites. KEGG-based pathway enrichment (via MetaboAnalyst) identified significant remodeling of arginine (Arg) and proline (Pro) metabolism, glycerophospholipid metabolism, Arg biosynthesis, linoleic acid metabolism, ether lipid metabolism, pyrimidine metabolism, valine/leucine/isoleucine biosynthesis, purine metabolism, glutathione metabolism, and cysteine (Cys) and methionine (Met) metabolism (*p* < 0.05), with β-alanine metabolism trending toward enrichment (*p* < 0.1) (Fig. 2J). These pathways delineate an integrated metabolic network coupling amino acid turnover, lipid remodeling, redox balance, and nitrogen management during extracellular protein catabolism. Complementary analysis using the SMPDB database (Fig. S5G) further supported these findings, revealing enrichment of phospholipid biosynthesis, aspartate metabolism, homocysteine degradation, and Met metabolism, consistent with engagement of transsulfuration-associated and redox-linked adaptations during extracellular protein catabolism.

At the metabolite level, extracellular protein catabolism induced coordinated remodeling across nitrogen-linked pathways, encompassing (i) Arg-cycle–linked nitrogen redistribution, (ii) acetylation-based nitrogen buffering, (iii) ammonia assimilation via the glutamate-glutamine axis, (iv) nucleotide turnover and β-alanine/CoA metabolism, and (v) transsulfuration and redox-linked detoxification. These processes were accompanied by remodeling of oxidative energy and NADPH-generating pathways that support redox balance under elevated nitrogen load (Fig. 2K, Supplementary Table S3).

Depletion of argininosuccinate and Arg, accompanied by accumulation of ornithine and N-acetyl-arginine indicated the engagement of partial Arg-cycle reactions that redistribute, rather than fully eliminate, nitrogen [49–51]. Given the lack of complete urea cycle in non-hepatic cells [30], elevated ornithine is consistent with nitrogen incorporation via Glu-centered transamination reactions, linking ammonia assimilation to mitochondrial carbon flux through the Glu-αKG node [52].

Nitrogen rerouting extended into nucleotide metabolism, as pyrimidine intermediates such as orotate and cytidine (log₂FC = −0.95, q = 1.4 × 10⁻^4^) were decreased, while β-alanine, the terminal product of pyrimidine catabolism and a precursor of panthothenate/CoA synthesis, was significantly increased (Fig. 2K). Reduced pantothenate (log₂FC = −0.35, q = 6.9 × 10⁻⁵) levels further suggest increased CoA utilization during oxidative and anabolic metabolism [30,53]. Together with increased inosine and inosine 5′-monophosphate (IMP), these changes indicate enhanced nucleotide turnover, nitrogen buffering, and mitochondrial energy metabolism in response to protein-derived nitrogen influx.

Amino-acid profiling further revealed activation of ammonia assimilation via the Glu-Gln axis [30]. As depicted in figure 2K, Gln was significantly increased (log₂FC = 0.38, q = 0.036), Glu was similarly upregulated (log₂FC = 0.28, q = 0.017), and α-KG showed a modest upward trend (log₂FC = 0.33, p = 0.13). This pattern indicates coordinated activation of GDH1 and transaminase reactions that couple ammonia incorporation to mitochondrial anaplerosis via the TCA cycle.

In parallel, acetylation-lined buffering pathways were engaged, as reflected by increased N-acetylglutamate (1.36-fold, q = 0.054, p = 0.026) and accumulation of multiple N-acetylated amino acids. Concurrent depletion of cystathione and homocysteine, together with increased ophthalmate, suggests an activation of the transsulfuration pathway and glutathione-linked detoxification as part of a broader nitrogen-redox adaptation.

Beyond nitrogen handling, lipid and energy metabolism were markedly remodeled. Upregulation of multiple acylcarnitines (acetyl-, margaroyl-, and long-chain species) and choline-containing phospholipids indicated enhanced fatty-acid oxidation and acetyl-CoA turnover. Elevated 2-ketogluconate, reflecting oxidative pentose-phosphate pathway and NADPH production, alongside the previously described increased β-alanine, underscores a coordinated metabolic response coupling oxidative energy production, redox adaptation, and nitrogen handling during extracellular protein catabolism.

Collectively, these data demonstrate that AML cells catabolize extracellular protein to meet both anabolic and energetic demands. Albumin supplementation liberates amino acids and carbon skeletons that sustain mTORC1 signaling, mitochondrial respiration, and proliferation while simultaneously engaging distinct nitrogen-handling programs that incorporate, redistribute, and detoxify excess nitrogen. These findings establish extracellular protein catabolism as a key component of AML metabolic plasticity that fuels oxidative metabolism and increases the requirement for effective ammonia handling [30].

### Extracellular protein catabolism induces ammonia accumulation and engages central nitrogen-handling pathways in AML

The observed enhancement nitrogen-handling activity inherently increases amino acid turnover and nitrogen flux, raising the possibility that extracellular protein catabolism increases ammonia accumulation as a measurable spillover of sustained nitrogen processing, with the potential to impose regulatory constraints at higher levels. To test this, we quantified extracellular ammonia concentrations following albumin supplementation. In HEL cells cultured at 0.5 × 10⁶ cells/mL for 24 hours, BSA significantly increased ammonia concentrations in the culture supernatant under both high-glucose (2 g/L) and glucose-restricted (0.5 g/L) conditions (Fig. 3A). A similar increase in ammonia levels was observed in primary AML blasts derived from two patients at diagnosis after 24 hours of BSA feeding (Fig. 3B). These findings demonstrate that extracellular protein catabolism elevates ammonia production in AML cells.

**Figure 3:**
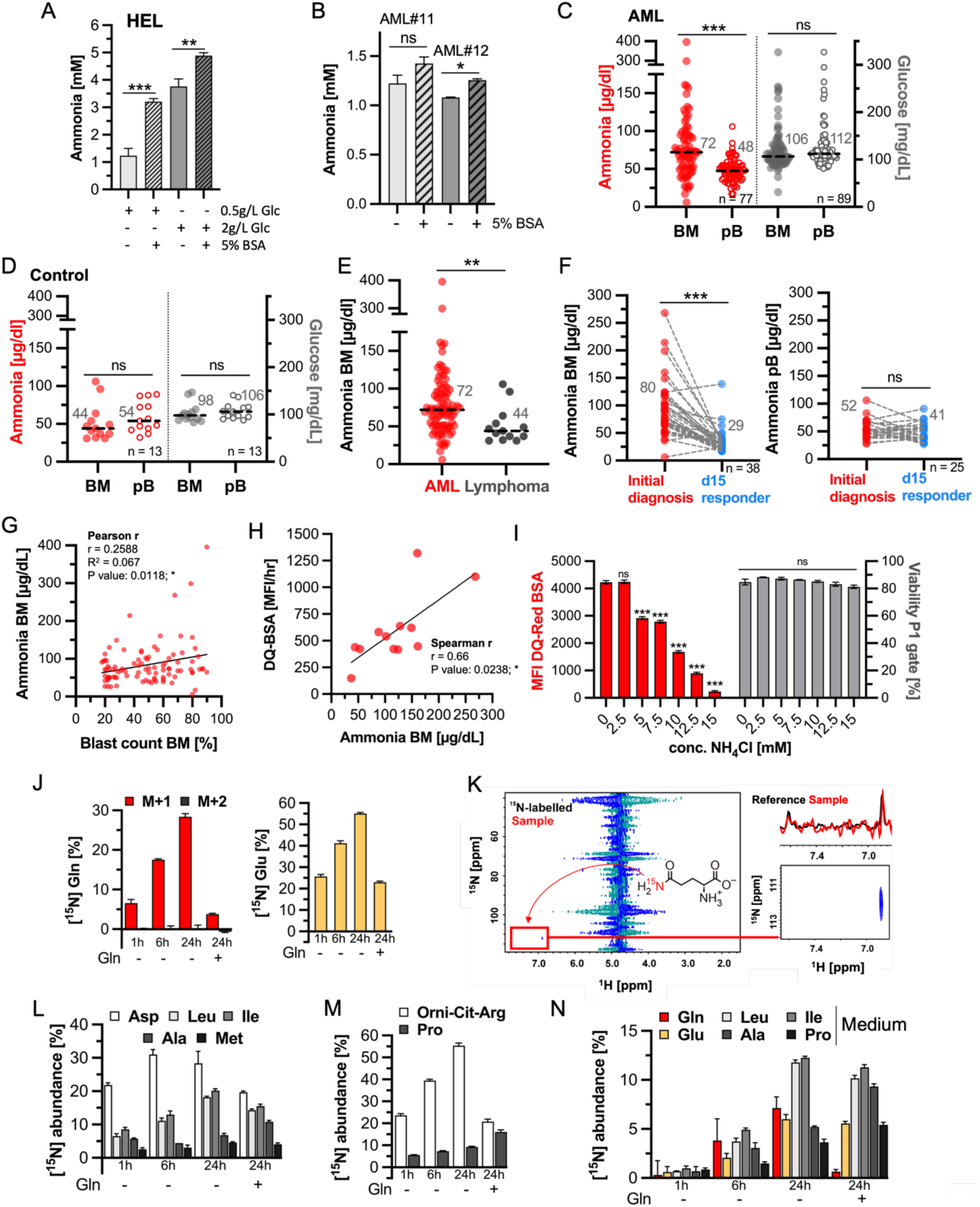
Extracellular protein catabolism induces ammonia accumulation and drives central nitrogen redistribution in AML. (A) Ammonia measurements (mmol/L) in supernatants of HEL cells cultured for 24 hours in medium containing standard glucose (Glc; 2 g/L) or reduced glucose levels (0.5 g/L), with or without 5% BSA (w/v). Unpaired t-test within each glucose condition; ***, p < 0.001. (B) Ammonia measurements (mmol/L) in supernatants of primary AML cells cultured for 24 hours in medium with or without 5% BSA (w/V) in duplicates. Unpaired t-test with Welch’s correction; *, p < 0.05. (C) Scatter dot plot of ammonia (red, µg/dL) and glucose (grey, mg/dL) concentrations in bone marrow (BM) and peripheral blood (PB) from 77 and 89 AML patients at first diagnosis, respectively. Lines and numbers indicate the median concentration of the patient cohort. Paired t-tests; ***, p < 0.001; ns, not significant. (D) Scatter dot plot of ammonia (red, µg/dL) and glucose (grey, mg/dL) concentrations in BM and PB from 13 lymphoma patients without bone marrow infiltration and with preserved normal hematopoiesis (normal BM controls). Lines and numbers indicate the median concentration of the patient cohort. Paired t-tests. (E) Scatter dot plot comparing BM ammonia concentrations (µg/dL) between AML patients and normal BM control patients. Lines and numbers indicate the median concentration of the patient cohorts. Unpaired t-test with Welch’s correction; **, p < 0.01. (F) Matched BM ammonia (left) and PB ammonia (right) concentrations (µg/dL) in 38 and 25 AML patients, respectively, at first diagnosis and during hematological complete remission (hCR; day 15 responders). Paired t-tests; ***, p < 0.001. (G) Pearson correlation between BM ammonia concentration (µg/dL) and BM blast count (%). (H) Spearman correlation between *ex vivo* protein catabolism rate (DQ-Red BSA; MFI/h) and BM ammonia concentration (µg/dL) in 10 AML patients at first diagnosis. (I) MFI of DQ-Red BSA (red bars) and cell viability (% in P1 gate) in HEL cells pretreated for 24 hours with 0, 2.5, 5, 7.5, 10, 12.5, or 15 mM NH₄Cl, followed by DQ-Red BSA exposure for 3 hours. Bar graphs represent mean ± SEM. Two-way ANOVA with Bonferroni’s post hoc test; ***, p < 0.001. (J) GC–MS analysis of HEL cells treated for 24 hours with 5 mM ¹⁵NH₄Cl. Bar graphs indicate mean relative abundance ± SD of ¹⁵N-glutamine (Gln; left) and ¹⁵N-glutamate (Glu; right) in cells cultured in glutamine-free medium for 1, 6, and 24 hours, and in standard RPMI medium for 24 hours. Red bars indicate Gln labeled with a single ¹⁵N atom (M+1); black bars indicate Gln labeled with two ¹⁵N atoms (M+2). (K) ¹H–¹⁵N heteronuclear single quantum coherence (HSQC) NMR spectrum of HEL cells treated for 24 hours with 5 mM ¹⁵NH₄Cl. (L) GC–MS analysis of HEL cells treated for 24 hours with 5 mM ¹⁵NH₄Cl. Bar graphs indicate mean relative abundance ± SD of transamination-derived ¹⁵N-aspartate (Asp), ¹⁵N-leucine (Leu), ¹⁵N-isoleucine (Ile), ¹⁵N-alanine (Ala), and ¹⁵N-methionine (Met) in cells cultured in glutamine-free medium for 1, 6, and 24 hours, and in standard RPMI medium for 24 hours. (M) GC–MS analysis of HEL cells treated for 24 hours with 5 mM ¹⁵NH₄Cl. Bar graphs indicate mean relative abundance ± SD of ¹⁵N-ornithine–citrulline–arginine (Orn–Cit–Arg) and ¹⁵N-proline (Pro) in cells cultured in glutamine-free medium for 1, 6, and 24 hours, and in standard RPMI medium for 24 hours. (N) GC–MS analysis of HEL cells treated for 24 hours with 5 mM ¹⁵NH₄Cl. Bar graphs indicate mean relative abundance ± SD of ¹⁵N-labeled Gln, Glu, Leu, Ile, Ala, and Pro in the culture medium from cells grown in glutamine-free medium for 1, 6, and 24 hours, and in standard RPMI medium for 24 hours.

To explore whether protein catabolism contributes to systemic metabolic changes in patients, we analyzed ammonia levels in the bone marrow (BM) and the peripheral blood (PB) in a cohort of 95 newly diagnosed AML patients (54 male, median age 63; 41 female, median age 64). The median BM ammonia concentration was 72 µg/dL (Fig. S6A), exceeding the normal serum reference range (11-60 µg/dL) in most cases. In 77 patients with matched PB samples, ammonia concentrations were significantly higher in BM (median 72 µg/dL) than in PB (median 48 µg/dL) (Fig. 3C). To control for potential sampling artifacts, glucose levels were measured in 89 patients. In contrast to ammonia, glucose levels were not significantly different between BM and PB (median 106 vs. 112 mg/dL, respectively), and remained within the normal range for non-fasting samples. These findings indicate that elevated ammonia in AML reflects active local production within the BM microenvironment rather than systemic variation or pre-analytical bias.

To determine whether elevated BM ammonia levels are specific to AML rather than a general feature of hematologic disease, we analyzed BM and PB ammonia concentrations in 13 lymphoma patients without BM infiltration and with preserved normal hematopoiesis (“healthy” control). In contrast to AML patients, these controls showed no significant difference in ammonia concentration between BM and PB (median 44 µg/dL vs. 54 µg/dL; Fig. 3D). Glucose levels were also similar in BM and PB (median 98 mg/dL vs. 106 mg/dL), supporting the notion that the observed ammonia gradient in AML is not due to sampling artifact or general BM physiology. We then directly compared BM ammonia concentrations between AML patients and lymphoma controls. Direct comparison of BM ammonia between cohorts revealed significantly higher levels in AML, confirming that local ammonia accumulation represents a disease-specific metabolic feature of AML (Fig. 3E).

We next asked whether BM ammonia levels are linked to leukemic burden. In 38 patients with matched BM plasma samples at diagnosis and day 15 of induction therapy, a timepoint at which responders typically show marked blast clearance. Median BM ammonia levels declined significantly from 80 µg/dL at diagnosis to 29 µg/dL on day 15 (empty BM, Fig. 3F, left). In contrast, no significant difference was observed in paired PB samples from 25 patients, where median ammonia concentrations changed from 52 µg/dL to 41 µg/dL over the same period (Fig. 3F, right). These findings reinforce that ammonia accumulation is a BM-restricted and disease-associated phenomenon in AML, closely tied to leukemic cell burden.

Since these findings indicate, that ammonia accumulation reflects active extracellular protein catabolism in AML, we asked whether ammonia itself feeds back on proteocatabolic activity. As depicted in figure 3I, exposure of HEL cells to increasing concentrations of ammonium chloride (NH_4_Cl) resulted in a graded reduction of DQ-Red BSA processing, indicating that elevated ammonia levels restrain extracellular protein catabolism in a concentration-dependent manner. This effect was observed at concentrations that precede overt loss of viability, suggesting that ammonia directly constrains proteocatabolic activity rather than merely reflecting nonspecific toxicity (Fig. 3I).

Given the observed accumulation of ammonia in the leukemic BM microenvironment, we next sought to define how AML cells detoxify and repurpose ammonia through central nitrogen metabolism. At physiological pH, ammonia (NH₃) exists primarily in its protonated moiety, ammonium (NH₄⁺), in both plasma and the cytoplasm (Fig. S6B) [55]. Intracellular assimilation of NH₄⁺ occurs through two complementary enzymatic routes: glutamate-ammonia ligase (or glutamine synthetase; GS, encoded by *GLUL*), which incorporates NH₄⁺ into Gln within the cytoplasm, and glutamate dehydrogenase 1 (GDH1, encoded by *GLUD1*), which catalyzes the reductive amination of α-KG to form Glu in the mitochondrial matrix. Although GDH1 is generally thought to catalyze oxidative deamination of Glu *in vivo*, it has been shown that high concentrations of α-KG and NH_4_^+^ can reverse this reaction, promoting Glu biosynthesis [56]. A third ammonia assimilation pathway, carbamoyl phosphate synthetase 1 (CPS1), initiates urea-cycle ammonia buffering but is largely restricted to hepatic mitochondria [30]. However, we detected CPS1 protein expression in our model AML cell line HEL (Fig. S6C), raising the possibility of cell line–specific carbamoyl-phosphate–linked reactions under protein scavenging conditions.

Thus, to trace the intracellular fate of NH_3_ in AML, HEL cells were treated with 5 mM ¹⁵N-labeled NH₄Cl, and intracellular labeling was analyzed by GC-MS over a 24-hour period. Incorporation of a single ¹⁵N label (M+1) into Gln increased progressively over time, reaching ∼28.4 ± 0.8% after 24 hours in Gln-deprived conditions (Fig. 3J, left). Parallel measurements showed robust ¹⁵N labeling of Glu, reaching ∼55.1 ± 0.4% in Gln-deprived conditions and 23.01+/-0.46% in the presence of Gln (Fig. 3J, right), consistent with direct NH_4_^+^ incorporation via GDH1. Notably, we did not detect M+2-labeled Gln, suggesting that GDH1-derived ^15^N-labeled Glu does not serve as a substrate for GS. Consequently, to confirm the atomic site of ^15^NH_4_^+^ incorporation in Gln we performed ¹H–¹⁵N HSQC-NMR. To distinguish between the two amino groups of Gln, we recorded reference spectra for both ^15^N-Glu and ^15^N-Gln and identified the chemical shift corresponding to the side chain amide group (δ-position) as a doublet at ^1^H6.85 ppm and ^15^N 112 ppm (Fig. S6D). The NMR spectrum from ¹⁵NH₄Cl-treated HEL cells confirmed exclusive ¹⁵N labeling at this side chain amide group (Fig. 3K), supporting GS-mediated NH_4_^+^ assimilation and excluding contribution of mitochondrial ¹⁵N-Glu to cytosolic Gln biosynthesis.

We next examined how GDH1-derived ¹⁵N-Glu contributes to downstream nitrogen distribution (Fig. S6B). GC-MS tracing revealed incorporation of ¹⁵N into multiple amino acids formed by transamination, including Asp, Ala, Leu, and Ile (Fig. 3L). In particular, Asp, produced from Glu by aspartate transaminase (AST), exhibited the highest labeling efficiency (∼31% at 6 h), highlighting the rapid redistribution of ammonia-derived nitrogen. Strikingly, ^15^N incorporation was also detected in proline (Pro) and in the ornithine–citrulline–arginine node (Fig. 3M), mirroring the elevations in ornithine, N-acetyl-arginine, and related intermediates observed in our global BSA-induced metabolomics dataset (Fig. 2K). This concordance reinforces that protein-derived nitrogen is funneled through the same Glu-linked pathways revealed by the tracer experiments. As depicted in figure S6B, these metabolites derive from Glu via the enzymes pyrroline-5-carboxylate synthase (P5CS), pyrroline-5-carboxylate reductase (P5CR) and ornithine aminotransferase (OAT), respectively. These findings indicate that ammonia-derived Glu is funneled into broader nitrogen metabolic pathways beyond direct elimination. Consistent with active nitrogen export, we detected ¹⁵N-labeled amino acids, including Gln, Glu, Leu, Ile, Ala, and Pro, in the extracellular medium (Fig. 3N), indicating that AML cells not only detoxify excess ammonia intracellularly but also release nitrogen-rich metabolites.

Collectively, these data demonstrate that ammonia is assimilated through spatially distinct GS- and GDH1-dependent routes and subsequently redistributed via transamination networks into amino acid pools and extracellular nitrogen-containing metabolites.

### Glutamate-ammonia ligase is a central metabolic safeguard that supports protein catabolism and ammonia tolerance in AML

Our isotopic tracing experiments demonstrated that AML cells detoxify ammonia through spatially distinct GS- and GDH1-dependent routes that incorporate excess nitrogen into Gln, Glu, and downstream amino acids. To determine which of these detoxification mechanisms is of predominant relevance in the disease context, we next determined the expression of ammonia-assimilating enzymes by performing meta-analysis of transcriptomic and proteomic datasets comparing primary AML patient samples to healthy hematopoietic stem and progenitor cells (HSPCs) [41–43] (Fig. 4A-C, Fig. S7A-C). This comparison revealed that within these patient cohorts, the most consistent upregulation at both the transcriptome and proteome levels was observed for GS/*GLUL*. GDH1/*GLUD1* was also upregulated, albeit to a lesser extent on transcriptomic levels, while *CPS1* expression was only present in one proteomic dataset [39] without transcriptional upregulation in AML (Fig. 4A). To visualize GS protein expression across individual samples, we plotted raw protein intensities from the De Boer and Kramer datasets [39, 40]. Both datasets showed consistently higher GS expression in AML samples compared to healthy CD34⁺ peripheral blood stem cells (PBSCs) and other controls (Fig. 4B). Further, we examined *GLUL* transcript levels in greater detail using two independent probe sets (200648_s_at and 217202_s_at) from the MILE transcriptome dataset. *GLUL* was significantly upregulated across multiple AML subtypes—including t(15;17), inv(16), t(8;21), and MLL-rearranged—as well as in other myeloid neoplasms such as chronic myeloid leukemia (CML) and myelodysplastic syndrome (MDS), but not in lymphoid malignancies (Fig. 4C). To extend these findings to a broader panel of cancer models, we analyzed *GLUL* expression across 1,450 cell lines from 74 cancer types in the DepMap 22Q4 dataset. AML cell lines exhibited *GLUL* expression levels that were predominantly above the median across the entire cohort (Fig. S7B). When focusing specifically on hematologic malignancy–derived cell lines, *GLUL* expression was significantly higher in AML and myeloproliferative neoplasm (MPN) cell lines compared to B-lymphoblastic leukemia/lymphoma, Hodgkin lymphoma, and non-Hodgkin lymphoma cell lines (Fig. S7C). Together, these findings demonstrate that *GLUL* is transcriptionally and translationally upregulated in AML and other myeloid neoplasms, representing a conserved metabolic adaptation in myeloid neoplasms and the most consistently upregulated ammonia-assimilating enzyme in AML.

**Figure 4:**
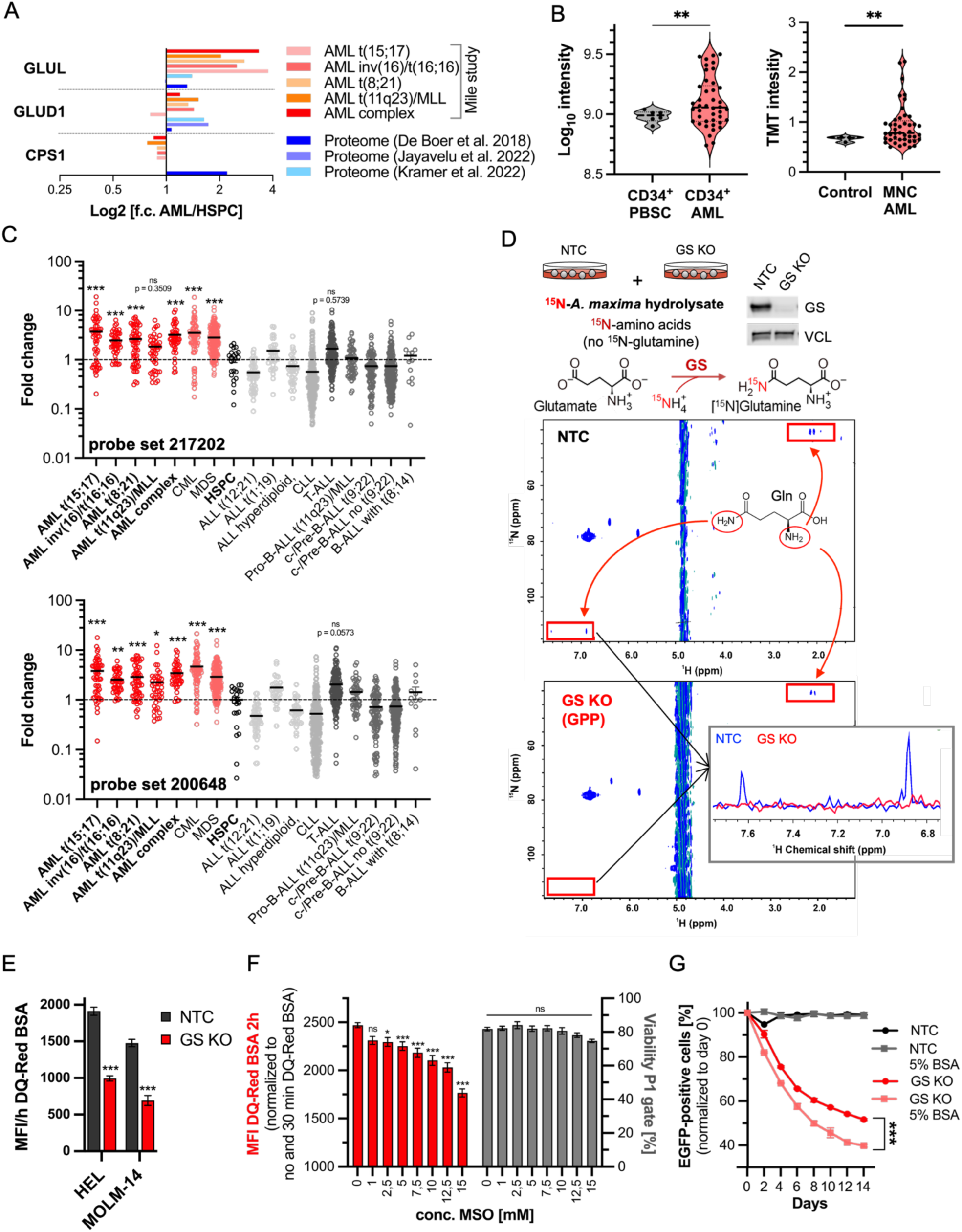
Glutamate–ammonia ligase links ammonia assimilation to extracellular protein catabolism in AML. (A) Integration of primary patient transcriptomic data from the MILE study and proteomic datasets [39–41] for enzymes and transporters involved in cellular ammonia assimilation. Fold changes are shown based on log₂-transformed or log₁₀-transformed expression values (log₁₀ for [39]), relative to normal hematopoietic stem and progenitor cells (HSPCs). (B) GS protein abundance in primary patient samples (CD34⁺ cells or mononuclear cells [MNCs]) compared to normal CD34⁺ peripheral blood stem cells (PBSCs) or healthy controls. Left: De Boer et al. (2018) [39]; right: Kramer et al. (2022) [40]. Welch’s t-test; **, p < 0.01. (C) GLUL mRNA abundance in primary patient samples (MILE study, probe set 217202 and 200648) compared with normal HSPCs. One-way ANOVA with Bonferroni’s post hoc test; *, p < 0.05; **, p < 0.01; ***, p < 0.001. (D) ¹H–¹⁵N heteronuclear single quantum coherence (HSQC) NMR spectra of HEL non-target control (NTC) and GS knockout (KO) cells after 24 hours of culture in the presence of ¹⁵N-labeled amino acids derived from a ¹⁵N-labeled *Arthrospira maxima* hydrolysate, which does not contain glutamine. GS knockout efficiency was assessed by SDS–PAGE followed by Western blotting. Vinculin (VCL) served as loading control. (E) Mean fluorescence intensity per hour (MFI/h) of DQ-Red BSA processing in HEL NTC and GS KO cells, and in MOLM-14 NTC and GS KO cells. Bar graphs represent mean ± SEM. Unpaired t-tests; ***, p < 0.001. (F) MFI of DQ-Red BSA after 3 hours (red bars), normalized to conditions without DQ-Red BSA and to unspecific fluorescence, and cell viability measured by flow cytometry (grey bars) in cells pretreated for 24 hours with increasing concentrations (0, 1, 2.5, 5, 7.5, 10, 12.5, and 15 mM) of the GS inhibitor L-methionine sulfoximine (MSO). Bar graphs represent mean ± SEM. Two-way ANOVA with Bonferroni’s post hoc test; *, p < 0.05; ***, p < 0.001. (G) Competitive growth assay of HEL NTC (black/grey) and GS KO cells (red/pink) cultured in the absence or presence of 5% BSA. The percentage of GFP-positive cells was assessed by flow cytometry every second day for 14 days. Extra sum-of-squares F test to evaluate differences in nonlinear growth-curve fitting; ***, p < 0.001.

Because amino acid turnover is the dominant intracellular source of ammonia, and protein degradation substantially increases amino acid turnover and nitrogen burden, we next examined whether GS function is linked to extracellular protein catabolism. To do so, we first depleted GS protein by CRISPR/Cas9-mediated knockout. The most effective single-guide RNA (sgRNA) – GPP - efficiently depleted full-length GS protein and markedly impaired proliferation (Fig. S7D-G) and was therefore used in subsequent experiments. We then cultured HEL NTC and HEL GS KO cells in the presence of ^15^N-labeled amino acids obtained from an *Arthrospira maxima* hydrolysate. A technical advantage of using a hydrolysate is the absence of ^15^N-labeled Gln due to the heat instability of Gln. As shown by the ^1^H-^15^N heteronuclear NMR spectra in figure 4D, we detected robust incorporation of the ^15^N-label into Gln in control cells but not in GS-deficient cells, confirming that GS mediates direct assimilation of ammonia generated during amino acid turnover.

Furthermore, if GS buffers the ammonia burden imposed by proteocatabolism, its depletion should constrain extracellular protein catabolism itself. As depicted in figure 4E, GS KO in HEL and MOLM-14 cells exhibited a significantly reduced protein catabolism rate compared to NTC control cells. Consistently, pharmacologic GS inhibition by L-methionine sulfoximine (MSO) markedly reduced lysosomal DQ-Red BSA processing in a dose-dependent manner without affecting overall viability (Fig. 4F). To determine whether GS contributes functionally to protein-supported proliferation, we performed competitive growth assays in the presence of 5% BSA. In this setting, the previously observed proliferation defect of GS KO cells (Fig. S7F/G) was further exacerbated when albumin served as the main nutrient source, highlighting the importance of GS when cells rely on extracellular protein catabolism (Fig. 4G).

Functionally, GS catalyzes the ATP-dependent synthesis of Gln from Glu and ammonia. A study, analyzing GS-function in pancreatic ductal adenocarcinoma (PDAC), showed that GS-mediated Gln synthesis is coupling α-KG-fueling of the TCA cycle with nitrogen anabolic processes, thereby supporting cancer cell growth and survival in a Gln-scarce microenvironment [31]. To test whether the observed growth defect of GS KO cells in AML could be attributed to impaired Gln production, we performed competitive assays across a range of extracellular Gln concentrations (600, 300, 100, 50 mg/L), whereas a Gln concentration of 300 mg/L corresponds to the standard Gln concentration in RPMI1640 medium (Fig. 5A). Even at a supraphysiological Gln concentrations of 600 mg/L, GS depletion resulted in a significant proliferation defect. The cultivation of GS KO cells in the presence of 50 mg/L Gln, a concentration that led to a loss of viability in control cells despite GS upregulation (Fig. S7H/I), causes an additional significant reduction of the GS KO cell population. This indicates that GS primarily stabilizes ammonia handling rather than serving as a Gln-supply enzyme under nutrient-replete conditions in AML.

**Figure 5:**
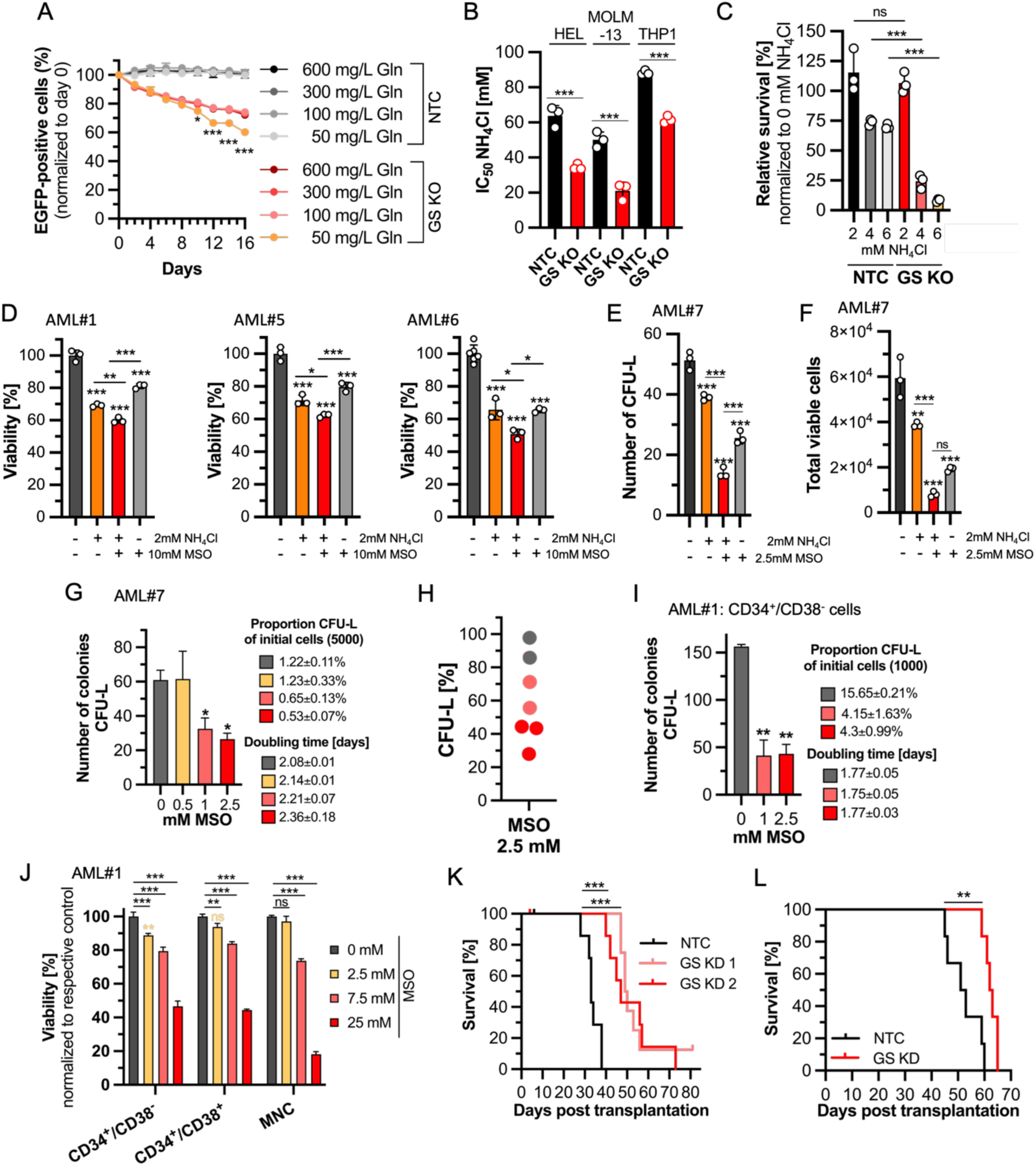
Glutamate–ammonia ligase is required for ammonia tolerance and leukemic growth in AML. (A) Glutamine (Gln) rescue experiment: competitive growth assay of HEL non-target control (NTC) and GS knockout (KO; sgRNA GPP) cells cultured for 16 days in the presence of 600, 300, 100, or 50 mg/L Gln. Points indicate the percentage of EGFP-positive cells. Error bars represent mean ± SD. Two-way ANOVA with Bonferroni’s post hoc test. Stars indicate significance between 50 mg/L Gln and 300 mg/L Gln in GS KO cells; *, p < 0.05; ***, p < 0.001. (B) IC₅₀ values of ammonium chloride (NH₄Cl) in HEL, MOLM-13, and THP-1 NTC and GS KO cells after 48 hours of treatment. Points indicate individual IC₅₀ values from biological replicates. Two-way ANOVA with Bonferroni’s post hoc test; ***, p < 0.001. (C) Viability assay following NH₄Cl treatment (2, 4, and 6 mM) in HEL NTC and GS KO cells for 48 hours, showing relative survival (%) normalized to the untreated control. Two-way ANOVA with Bonferroni’s post hoc test; *, p < 0.05; **, p < 0.01. (D) Viability (%) of three primary AML samples (AML#1, AML#6, AML#7) treated with 10 mM L-methionine sulfoximine (MSO) and/or 2 mM NH₄Cl for 48 hours. Stars above bars indicate significance compared to the untreated control. One-way ANOVA with Bonferroni’s post hoc test; *, p < 0.05; **, p < 0.01; ***, p < 0.001. (E) Colony formation assay showing the number of CFU-L colonies formed by primary AML blasts (AML#8) treated with 2.5 mM MSO and/or 2 mM NH₄Cl. Cells were cultured for 21 days under hypoxic conditions. (F) Colony formation assay showing the total number of viable cells derived from primary AML blasts (AML#8) treated with 2.5 mM MSO and/or 2 mM NH₄Cl after 21 days of culture under hypoxia. (G) Colony formation assay of primary AML blasts treated with 0, 0.5, 1, or 2.5 mM MSO and cultured for 21 days under hypoxia. Left: number of CFU-L colonies (bar graphs). Proportion of colony-forming units (CFU) relative to initially seeded cells and doubling time are indicated in the legends. (H) Summary of CFU assays shown in Supplementary Fig. S8. Viability was normalized to the respective control samples. *Ex vivo* responders are shown in red (0–50% viability upon MSO treatment) and light red (50–75% viability upon MSO treatment). (I) Colony formation assay of CD45^dim^/CD34⁺/CD38⁻ leukemia-initiating cells (LICs; AML#1) treated with 0, 1, or 2.5 mM MSO and cultured for 21 days under hypoxia. Left: number of CFU-L colonies (bar graphs). Proportion of CFU relative to initially seeded cells and doubling time are indicated in the legends. Bar graphs represent mean ± SD of two technical replicates. One-way ANOVA with Bonferroni’s post hoc test; *, p < 0.05; **, p < 0.01. (J) Relative viability (%) of CD45^dim^/CD34⁺/CD38⁻ LICs, CD45^dim^/CD34⁺/CD38⁺ progenitor cells, and mononuclear cells (MNCs) from AML#1 treated with 0, 2.5, 7.5, or 25 mM MSO for 48 hours under hypoxia. Bar graphs represent mean ± SD of three technical replicates. Two-way ANOVA with Bonferroni’s post hoc test; **, p < 0.01; ***, p < 0.001. (K) Xenotransplantation of THP-1 cells with GS knockdown (KD) showing probability of survival. Mantel–Cox test. Median survival: NTC = 33 days; GS KD 1 = 49.5 days; GS KD 2 = 47 days; ***, p < 0.001. (L) Transplantation of MN1 cells (NTC and GS KD sh5) into non-lethally irradiated C57BL/6J mice, showing probability of survival. Mantel–Cox test. Median survival: NTC = 52 days; GS KD = 62 days; **, p < 0.01.

To determine whether the principal function of GS in AML is ammonia detoxification, we next tested if GS KO cells exhibited an increased sensitivity to exogenous NH_4_Cl treatment. In three distinct AML cell lines the loss of GS significantly increased the sensitivity of the cells to ammonia (Fig. 5B). Similarly, increasing concentrations of NH_4_Cl (2, 4, 6 mM) led to a higher toxicity in GS KO compared with NTC cells (Fig. 5C). To validate this finding in primary AML, we exposed patient-derived blasts to ammonium and the previously used GS inhibitor MSO. In three independent AML samples (AML#1, AML#6, AML#7) NH_4_Cl supplementation strongly reduced cell viability and MSO treatment further sensitized the cells to ammonia stress (Fig. 5D). Similarly, colony formation (CFU-L) and the number of total viable cells were impaired upon NH_4_Cl treatment and this effect was amplified by GS inhibition (Fig. 5E/F). Together, these results demonstrate that GS activity is essential to buffer ammonia stress and sustain leukemic growth in both cell lines and primary AML blasts.

Given the role of GS in buffering ammonia stress, we next examined its functional importance in leukemogenesis in *ex vivo* primary AML blasts and in *in vivo* mouse models. The pharmacologic inhibition of GS using MSO suppressed colony formation of primary AML blasts in methylcellulose-based colony-forming unit (CFU-L) assays in a dose-dependent manner and increased the overall doubling time (Fig. 5G/H, Fig. S8A-F). GS inhibition also significantly reduced CFU-L numbers and viability from CD34⁺/CD38⁻ leukemia-initiating cells (LICs), indicating that GS supports both leukemic bulk and stem-like compartments (Fig. 5I/J). *In vivo*, shRNA-mediated GS knockdown in THP-1 and MN1-driven leukemias significantly delayed disease progression and extended overall survival (15.25 days in NSG mice and 10 days in the MN1 model) (Fig. 5K/L, Fig. S8G-L). Together, these data identify GS as a recurrently upregulated and functionally required enzyme in AML.

Collectively, our findings demonstrate that GS is transcriptionally upregulated, functionally essential, and mechanistically required to detoxify ammonia generated during extracellular protein catabolism in AML. By incorporating ammonia into Gln, GS prevents intracellular nitrogen overload and stabilizes sustained amino acid and extracellular protein catabolism. This buffering capacity allows AML cells to maintain anabolic and oxidative metabolism under proteocatabolic conditions but also exposes a critical dependence on GS activity to buffer the ammonia externality generated by their protein-scavenging strategy. Thus, GS represents a central metabolic node and a potential therapeutic vulnerability in AML.

## 4. Discussion

### Proteocatabolism establishes a regulated nitrogen regimen in AML

Macropinocytosis and lysosomal proteocatabolism are well-established nutrient-scavenging strategies in cancer, particularly in KRAS-driven pancreatic tumors, where extracellular protein supplies amino acids to sustain growth under nutrient limitation [21–24]. Our data extend this framework to AML and reveal an additional, underappreciated dimension: extracellular protein catabolism provides a substantial source of bioavailable nitrogen that is efficiently incorporated into cellular metabolism rather than immediately eliminated. This observation aligns with broader concepts in cancer nitrogen metabolism, which emphasize coordinated nitrogen acquisition and redistribution to support anabolic demand and metabolic flexibility [28,30].

AML blasts, particularly those with monocytic differentiation, exhibit elevated rates of extracellular protein catabolism compared with other hematologic malignancies and normal hematopoietic populations, indicating that proteocatabolism represents a disease-enriched metabolic adaptation rather than a generic feature of proliferation (Fig. 1; Fig. S2; Fig. S3). Uptake of high–molecular weight dextran and its correlation with DQ-BSA degradation across AML cell lines indicate that bulk uptake contributes substantially to extracellular protein acquisition (Fig. 1I/J; Fig. S4). Partial inhibition by EIPA supports a major, though not exclusive, role for macropinocytosis in establishing proteocatabolic capacity (Fig. 1K; Fig. S4). Functionally, protein-derived nutrients are productively integrated into anabolic signaling and mitochondrial metabolism, supporting proliferation even under conditions of amino acid or glucose limitation (Fig. 2A–I; Fig. S5).

Importantly, the metabolic contribution of proteocatabolism extends beyond carbon metabolism. Metabolomic and isotope-tracing analyses demonstrate that protein-derived nitrogen is actively redistributed through central nitrogen-handling nodes, including the Glu-Gln axis selective Arg-cycle–linked reactions, and nucleotide-associated metabolic processes (Fig. 2J/K; Fig. 3J–N). Together, these data support a model in which extracellular protein catabolism establishes a high-throughput proteocatabolic nitrogen regimen in AML, conferring metabolic flexibility through enhanced nitrogen availability and routing capacity.

### Ammonia spillover reflects high nitrogen flux and imposes regulatory constraints

Within this framework, the ammonia accumulation observed during proteocatabolism *in vitro* and in the BM of AML patients reflects the spillover of reduced nitrogen under conditions of sustained nitrogen processing, rather than a primary metabolic defect. BM ammonia concentrations are elevated in AML compared with paired PB and decline with effective cytoreductive therapy, indicating that ammonia accumulation is a local and disease-linked phenomenon (Fig. 3C–F). Across primary samples, *ex vivo* DQ-Red BSA processing correlates with ammonia concentrations measured in matched BM aspirates (Fig 3H), further supporting a direct link between proteocatabolic throughput and nitrogen spillover.

Although AML cells tolerate ammonia concentrations within this biologically relevant range, ammonia is not inert. *Ex vivo* and *in vitro* exposure to increasing ammonia concentrations leads to graded impairment of proliferation, viability, and clonogenic capacity (Fig. 5B-F), and ammonia also restrains proteocatabolic throughput itself (Fig. 3I). Thus, ammonia functions as a regulatory externality of the nitrogen regimen, imposing constraints that must be actively managed to preserve metabolic homeostasis and sustain proteocatabolic growth.

### GS functions as a stabilizer of the proteocatabolic nitrogen regimen

We next sought to define how AML cells buffer ammonia to remain within a viable operating range. Stable isotope tracing identified both glutamate–ammonia ligase (GS) and glutamate dehydrogenase (GDH1) as routes of ammonia assimilation. However, analyses of transcriptomic and proteomic data from multiple AML patient cohorts consistently identified GS as the dominant upregulated enzyme, whereas GDH1 showed more modest induction and CPS1 lacked consistent expression (Fig. 4A-C; Fig. S7A). This prominent role of GS is consistent with its emerging recognition as a central metabolic node in cancer, coupling ammonia assimilation to metabolic adaptation and tumor fitness [57]. This pattern extends to large-scale cancer cell line datasets, suggesting a lineage-associated adaptation among myeloid malignancies (Fig. S7B/C).

Functionally, GS activity is tightly coupled to proteocatabolic metabolism. Incorporation of ^15^N from amino acid hydrolysates into Gln is abolished in GS-deficient cells (Fig. 4D), establishing GS as a principal mediator of ammonia assimilation derived from extracellular protein breakdown. GS dependency is further amplified under proteocatabolic growth conditions when supplementation with albumin exacerbates the competitive growth disadvantage of GS-deficient cells, and pharmacologic or genetic GS inhibition reduces DQ-Red BSA processing at concentrations that do not impair viability (Fig. 4E-G). These findings indicate that GS not only buffers ammonia toxicity, similar to its function during erythropoiesis [58], but also prevents ammonia-mediated negative feedback on proteocatabolic activity, thereby stabilizing the nitrogen regimen.

Consistent with this role, genetic or pharmacologic GS inhibition impairs proliferation in AML cell lines, reduces colony formation and viability of primary AML blasts and leukemia-initiating cells (Fig. 5D-J), and delays leukemia progression *in vivo* (Fig. 5K/L). Importantly, excess extracellular Gln fails to rescue GS-deficient cells across a wide concentration range (Fig. 5A), indicating that GS is required primarily for ammonia assimilation rather than Gln supply. Thus, in contrast to nutrient-deprived solid tumors such as pancreatic ductal adenocarcinoma, where GS supports Gln synthesis to sustain anaplerosis under Gln scarcity [31,59], AML exhibits a distinct metabolic context in which nitrogen excess and ammonia buffering define the vulnerability.

### Conceptual implications

Together, our findings define a metabolic framework in which extracellular protein catabolism enables AML cells to establish a regulated, high-throughput nitrogen regimen. Proteocatabolism provides bulk access to reduced nitrogen that is actively routed into central metabolic pathways, supporting biosynthetic and energetic demands in the BM microenvironment. Ammonia accumulation emerges as a predictable spillover of sustained nitrogen throughput, creating a conditional liability that restrains proteocatabolic activity unless effectively buffered. GS functions as a regimen stabilizer, defining the upper limit of sustainable nitrogen handling capacity and enabling AML cells to exploit the benefits of proteocatabolism without incurring self-limiting toxicity. This work reframes proteocatabolism from a nutrient-scavenging adaptation into a lineage-compatible nitrogen management strategy with inherent and potentially exploitable vulnerabilities.

### Limitations and future directions

AML is a heterogeneous disease, and while our analyses span primary patient samples and multiple experimental models, they do not capture the full spectrum of genetic and differentiation states. Future work will be required to define which AML subtypes most strongly engage proteocatabolic nitrogen regimens and how this metabolic trait relates to clinical behavior. In addition, our pharmacological assessment of macropinocytosis relies on EIPA, which is not a selective macropinocytosis inhibitor. Recently developed, more selective pharmacological approaches to inhibit macropinocytosis [60] should help to further define the quantitative contribution of macropinocytosis relative to alternative protein-uptake pathways in AML. Our study focused primarily on cell-intrinsic nitrogen handling and did not directly interrogate how ammonia spillover influences non-malignant components of the BM niche, an important future direction given emerging evidence that tumor-derived ammonia can modulate immune and stromal cell function without inducing overt toxicity [61,62]. Finally, translating GS dependence into therapeutic strategies will require development of clinically tractable inhibitors and combination approaches that constrain both proteocatabolic flux and ammonia buffering capacity.

## Supplementary Materials

### Supplementary figures

**Supplementary figure S1:**
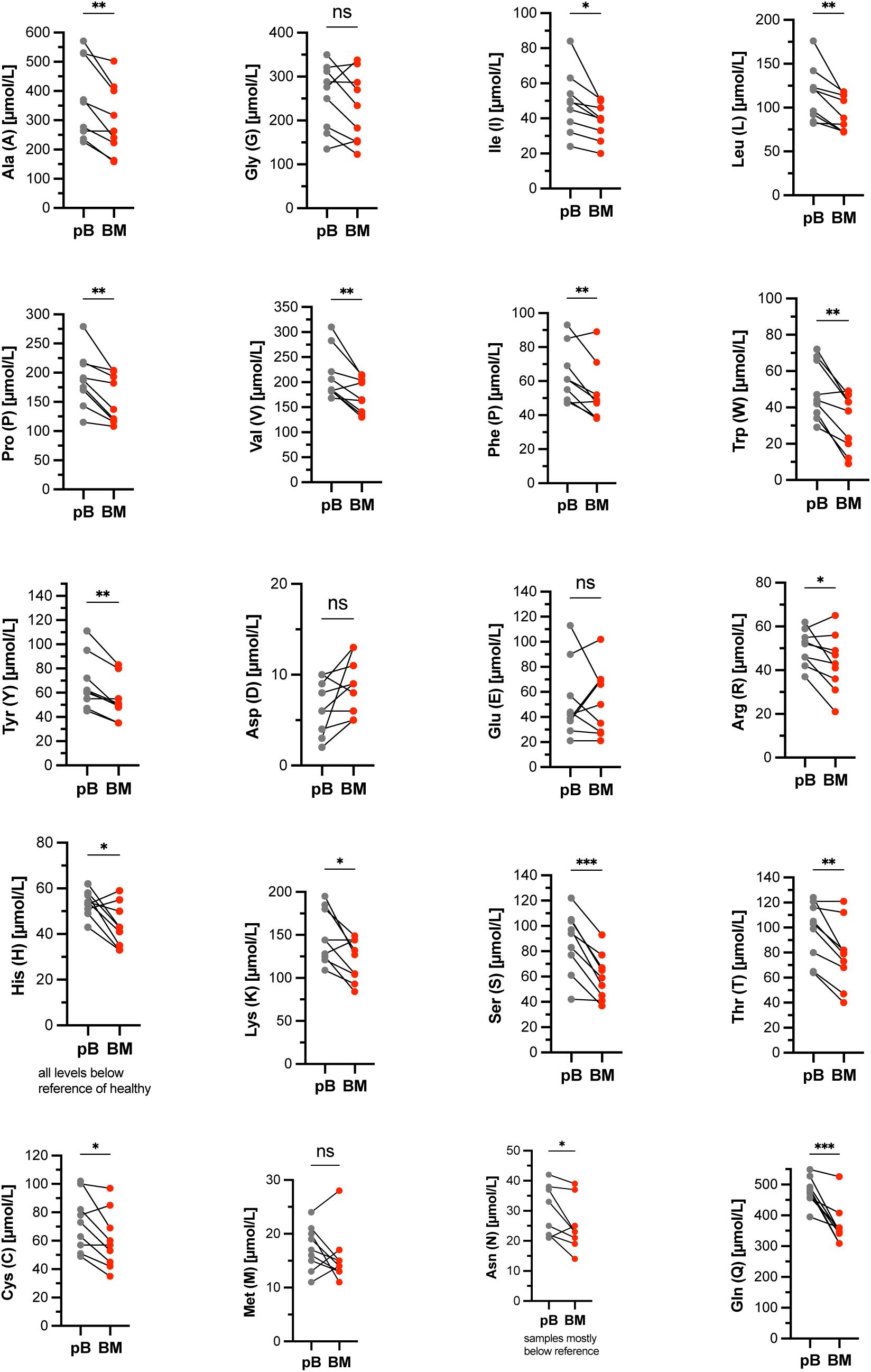
Free amino acid concentration in the peripheral blood and bone marrow of AML patients. Measurements of free amino acids in matched peripheral blood (PB) and bone marrow (BM) samples of seven AML patients at first diagnosis. Paired t-test; *, p <0.05; **, p <0.01; ***, p <0.001.

**Supplementary figure S2:**
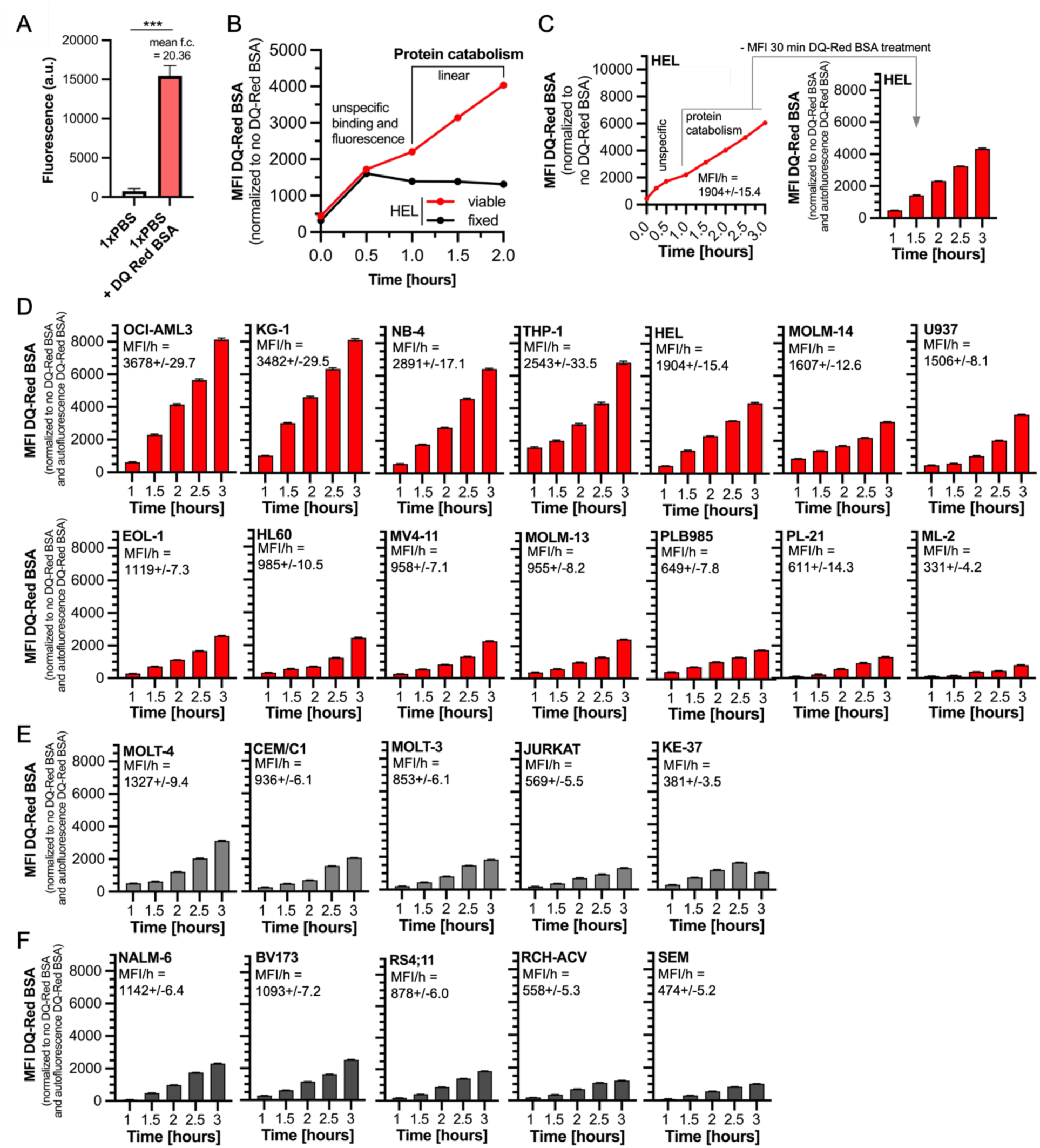
AML cells take up extracellular protein in form of serum albumin. (A) Plate-based measurement of DQ-Red BSA fluorescence (arbitrary units, a.u.). Bar graphs represent mean ± SEM. Unpaired t-test; ***, p < 0.001. (B) Mean fluorescence intensity (MFI) of DQ-Red BSA in viable (red) and fixed (black) HEL cells after 0.5, 1, 1.5, and 2 hours of treatment. Unspecific binding and fluorescence are observed up to 1 hour of treatment. The linear increase between 1 and 2 hours indicates lysosomal DQ-Red BSA processing (protein catabolism). (C) Measurement of MFI of DQ-Red BSA in HEL cells after 0, 0.15, 0.5, 1, 1.5, 2, 2.5, and 3 hours of treatment, normalized to the no DQ-Red BSA control. Within the linear range of the fluorescence signal between 1 and 3 hours, lysosomal conversion of DQ-Red BSA occurs. The slope of the resulting linear regression indicates the rate of protein catabolism expressed as MFI/h (left). The MFI value corresponding to unspecific fluorescence resulting from autofluorescence of DQ-Red BSA without lysosomal processing at 0.5 hours was subtracted from subsequent values and is shown as a bar graph. Bar graphs represent mean ± SEM. (D) MFI of DQ-Red BSA after 1, 1.5, 2, 2.5, and 3 hours of treatment in 14 distinct AML cell lines. Bar graphs represent mean ± SEM. In addition, the protein catabolism rate calculated from the linear regression is shown as MFI/h ± SEM. (E) MFI of DQ-Red BSA after 1, 1.5, 2, 2.5, and 3 hours of treatment in five distinct T-ALL cell lines. Bar graphs represent mean ± SEM. In addition, the protein catabolism rate calculated from the linear regression is shown as MFI/h ± SEM. (F) MFI of DQ-Red BSA after 1, 1.5, 2, 2.5, and 3 hours of treatment in five distinct B-ALL cell lines. Bar graphs represent mean ± SEM. In addition, the protein catabolism rate calculated from the linear regression is shown as MFI/h ± SEM.

**Supplementary figure S3:**
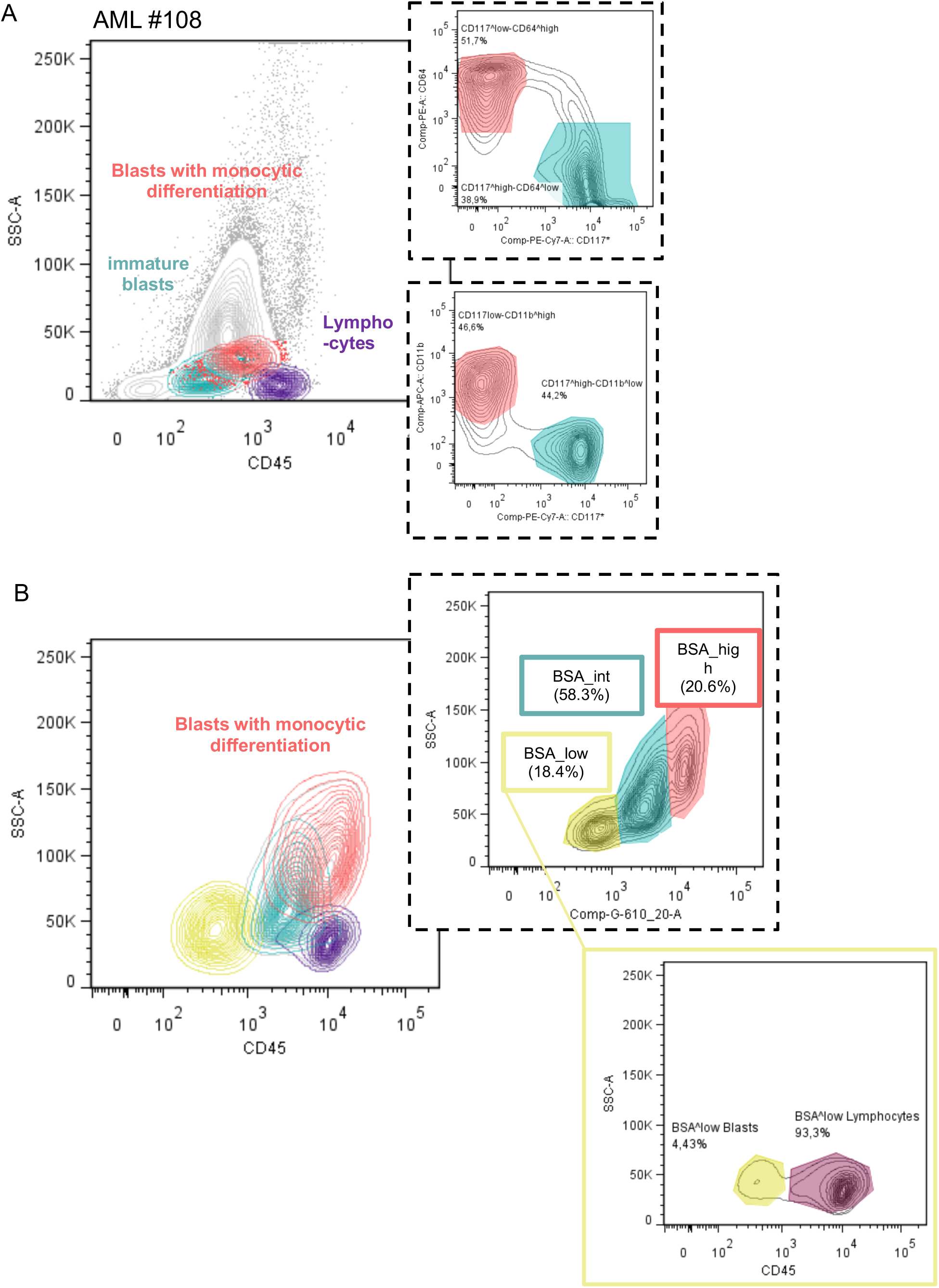
Extracellular protein catabolism is enriched in AML with monocytic differentiation features. (A) Representative flow cytometric immunophenotyping from clinical routine diagnostics of the primary AML sample AML12-25. (Left) CD45/SSC-A plot showing blasts with monocytic differentiation, immature blasts, and lymphocytes. (Right) Immunophenotypic differentiation of blasts based on CD117/CD64 expression (upper panel) and CD117/CD11b expression (lower panel), used to classify cells as monocytoid-differentiated or immature. (B) Flow cytometric analysis of a Ficoll-separated sample from the same patient (AML12-25). (Left) CD45/SSC-A plot color-coded according to DQ-BSA uptake. (Right, upper) DQ-Red BSA fluorescence plotted against SSC-A defining DQ-BSA_low, DQ-BSA_int, and DQ-BSA_high subpopulations. (Right, lower) Backgating of the DQ-BSA_low subpopulation into CD45/SSC-A space to distinguish lymphocytes from DQ-BSA_low blasts, highlighted by a yellow boxed gate.

**Supplementary figure S4:**
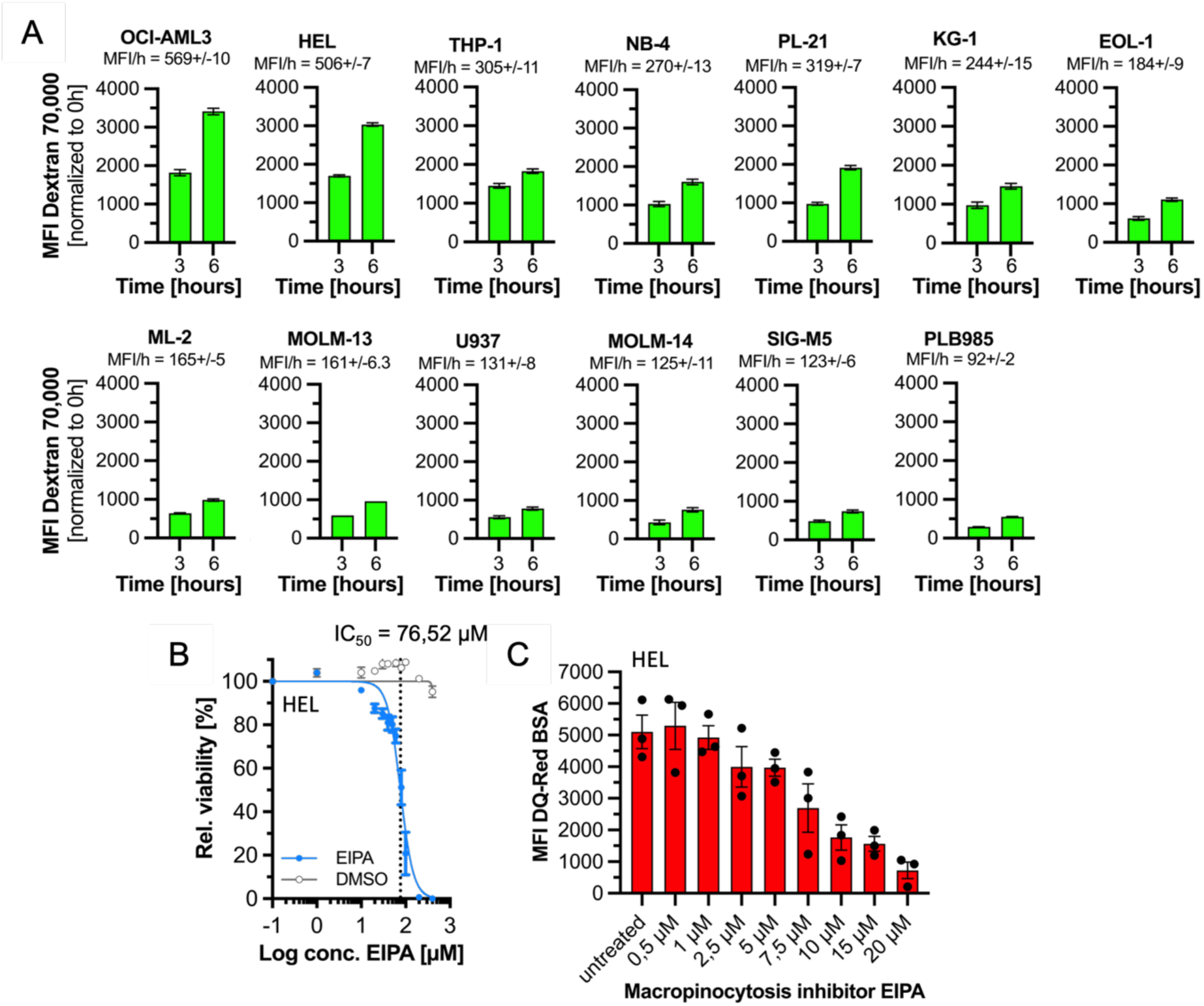
Dextran uptake assays and EIPA validation for macropinocytosis-dependent protein scavenging. (A) Mean fluorescence intensity (MFI) of dextran (70,000 MW) after 3 and 6 hours of treatment in 13 distinct AML cell lines. Bar graphs represent mean ± SEM. In addition, the macropinocytosis rate calculated from the linear regression is shown as MFI/h ± SEM. (B) ATP-based cell viability assay of HEL cells treated for 24 hours with increasing concentrations of EIPA (5-(N-ethyl-N-isopropyl)amiloride) or corresponding DMSO concentrations (solvent control). The IC₅₀ value of EIPA in HEL cells is 76.52 µM. (C) Mean fluorescence intensity (MFI) of DQ-Red BSA in HEL cells treated for 24 hours with increasing concentrations of EIPA.

**Supplementary figure S5:**
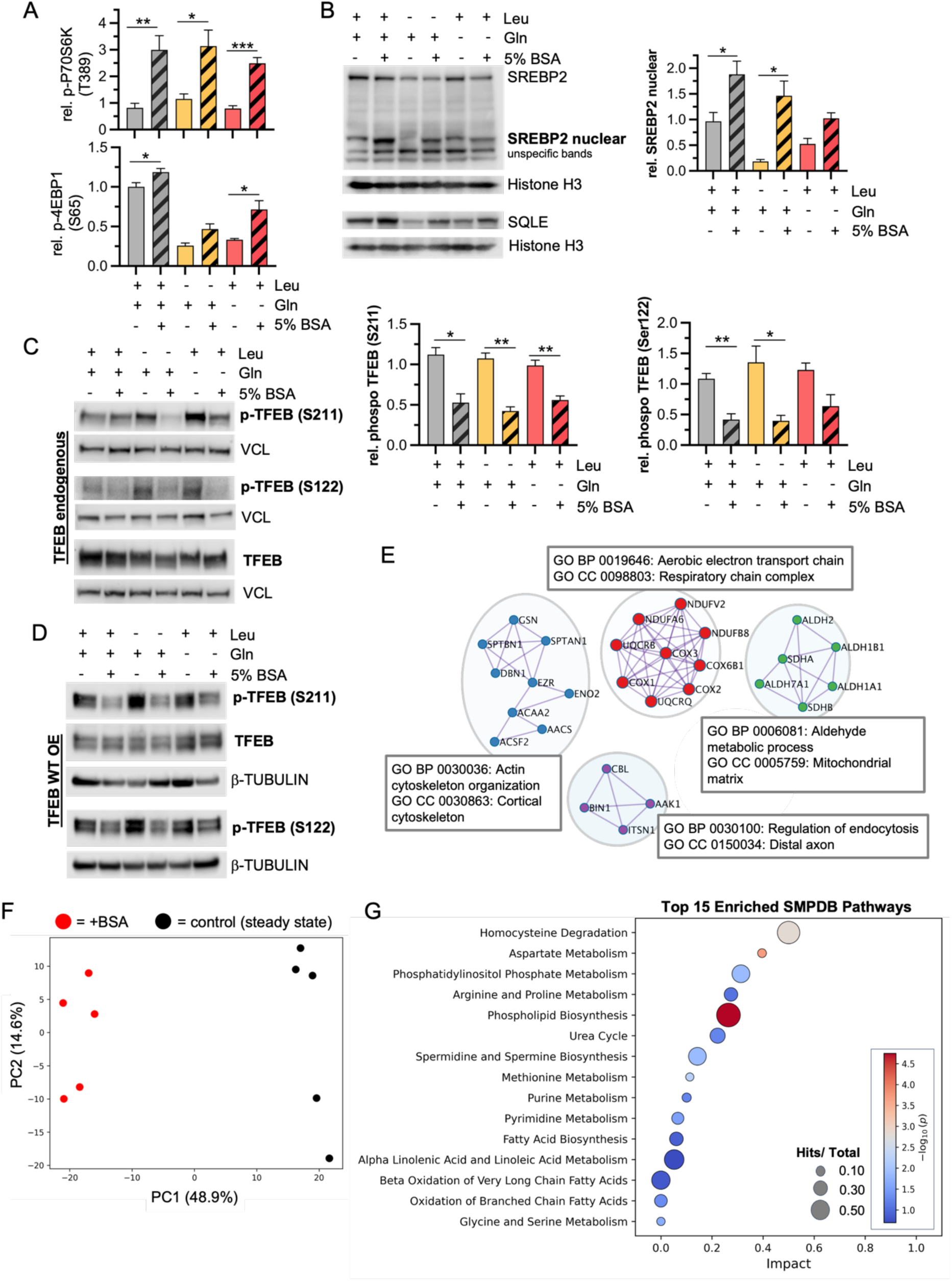
Extracellular albumin supplementation activates anabolic signaling and drives broad metabolic remodeling in AML cells. (A) Quantification of phospho-P70S6K (T389) and phospho-4EBP1 (S65) from three independent experiments of HEL cells cultured for 24 hours in full, leucine-deprived (−Leu), or glutamine-deprived (−Gln) medium, with or without 5% BSA (w/v). (B) Western blot (left) and quantification (right; SREBP2 only) of HEL cells cultured for 24 hours in full, −Leu, or −Gln medium, with or without 5% BSA (w/v). Detection of endogenous SREBP2 and squalene epoxidase (SQLE). Histone H3 served as loading control. (C) Western blot (left) and quantification (right) of HEL cells cultured for 24 hours in full, −Leu, or −Gln medium, with or without 5% BSA (w/v). Detection of endogenous phospho-TFEB (S211), phospho-TFEB (S122), and total TFEB. Vinculin (VCL) served as loading control. (D) Western blot analysis of HEL cells overexpressing wild-type TFEB and cultured for 24 hours in full, −Leu, or −Gln medium, with or without 5% BSA (w/v). Detection of phospho-TFEB (S211), phospho-TFEB (S122), and total TFEB. β-tubulin served as loading control. (E) Protein interaction network derived from the global proteomics dataset of HEL cells cultured for 7 days with 5% BSA (w/v), showing clusters identified in pathway enrichment analyses based on GO biological processes (BP) and GO cellular components (CC). (F) Principal component analysis (PCA) of the global metabolomics dataset of HEL cells treated for 24 hours with 5% BSA (w/v). (G) Pathway enrichment analysis of the global metabolomics dataset of HEL cells cultured for 24 hours with 5% BSA (w/v), showing the top 15 enriched SMPDB pathways. Bubble size represents the metabolite hit/total ratio. Red–blue color scale represents −log₁₀(p-value).

**Supplementary figure S6:**
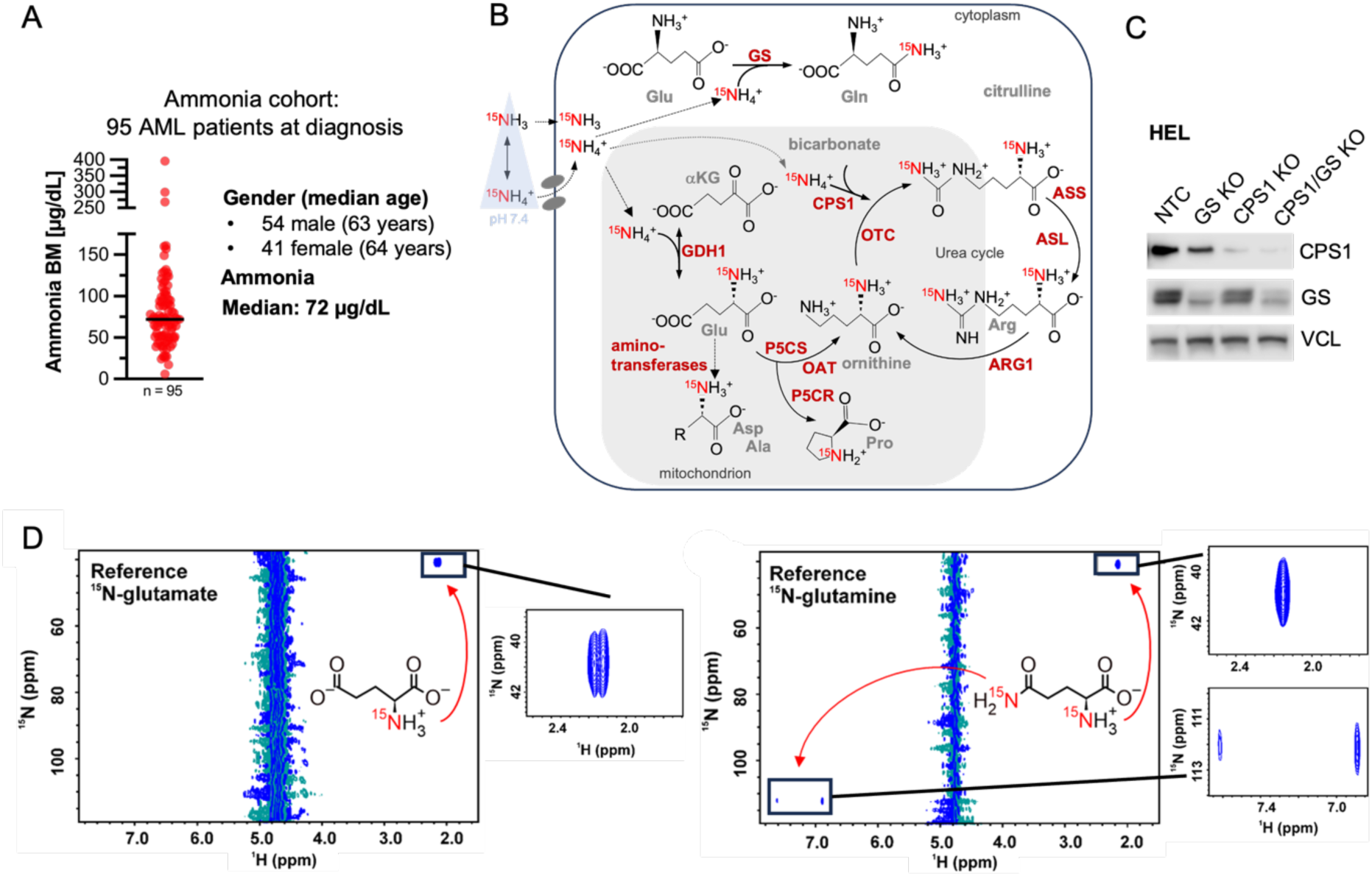
Bone marrow ammonia distributions and supporting data for ammonia assimilation and tracing experiments. (A) Scatter dot plot showing the distribution of bone marrow (BM) ammonia concentrations measured in 95 AML patients at first diagnosis. The median ammonia concentration, as well as patient gender distribution and median age, are indicated. (B) Schematic representation of ¹⁵N-ammonium detoxification at physiological pH (7.4). At physiological pH, ammonia is predominantly present in its ionized form, ammonium (NH₄⁺). In the cytoplasm, glutamate–ammonia ligase (glutamine synthetase; GS) catalyzes the conversion of glutamate and ammonium to glutamine, resulting in labeling of the amide nitrogen at the δ position. In mitochondria, detoxification of ¹⁵N-ammonium occurs through conversion of α-ketoglutarate (α-KG) to glutamate by glutamate dehydrogenase 1 (GDH1). An additional ammonium detoxification pathway, primarily described in the liver, involves conversion by carbamoyl phosphate synthetase 1 (CPS1) and subsequent entry into the urea cycle. (C) Western blot analysis detecting endogenous levels of glutamate–ammonia ligase (GS) and carbamoyl phosphate synthetase 1 (CPS1). Vinculin (VCL) served as loading control. (D) ¹H–¹⁵N heteronuclear single quantum coherence (HSQC) NMR reference spectra of ¹⁵N-glutamate and ¹⁵N-glutamine (left).

**Supplementary figure S7:**
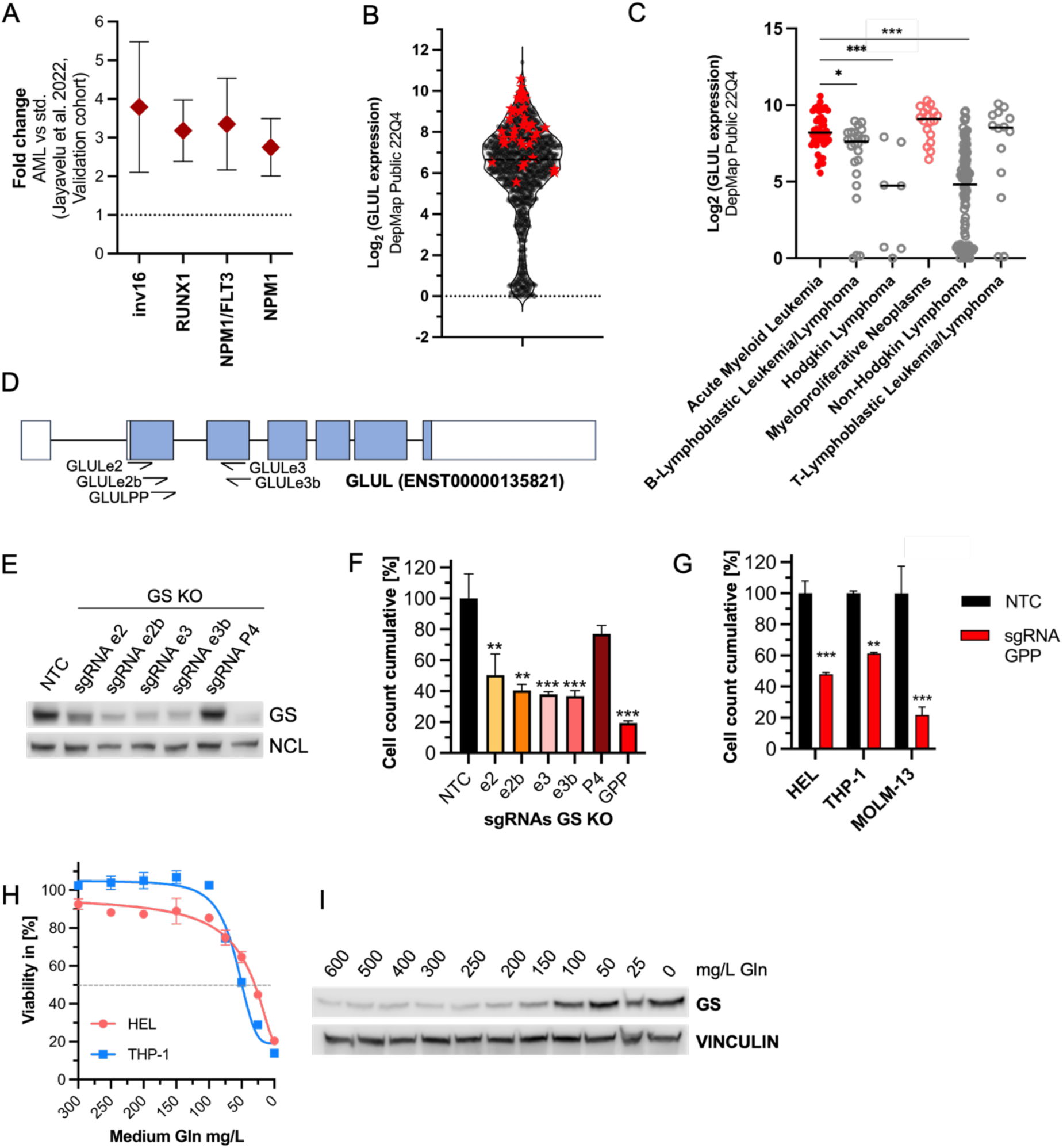
GS expression across cancer models and validation of GS genetic perturbation in AML cells. (A) Mean ± SEM of GS protein abundance in primary patient samples (mononuclear cells, MNCs) compared to an AML cell line standard (std.) [41]. (B) Sorted expression of *GLUL* across cancer cell lines from the DepMap Public 22Q4 dataset. AML cell lines are highlighted as red stars. (C) Log₂-transformed *GLUL* expression across cell lines from the DepMap Public 22Q4 dataset, grouped by primary hematologic disease. One-way ANOVA with Bonferroni’s post hoc test; *, p < 0.01; ***, p < 0.001. (D) Schematic representation of the GLUL gene locus (ENST00000135821), indicating the positions of the single-guide RNAs (sgRNAs) used for GS targeting. (E) Western blot analysis of HEL cells depleted (knockout) of GS using six distinct sgRNAs. Nucleolin (NCL) served as loading control. e = exon; P4 = GLUL pseudogene; GPP = sgRNA targeting all GLUL gene copies, including pseudogenes. (F) Cumulative cell counts on day 15 in HEL cells depleted (knockout) of GS using six distinct sgRNAs. Error bars represent mean ± SEM. One-way ANOVA with Bonferroni’s post hoc test performed on day 15; **, p < 0.01; ***, p < 0.001. (G) Cumulative growth (%) of HEL, THP-1, and MOLM-13 cells depleted of GS using sgRNA GPP, assessed on day 7. Two-way ANOVA with Bonferroni’s post hoc test; **, p < 0.01; ***, p < 0.001. (H) Viability assay measured by relative ATP production in HEL and THP-1 cells cultured for 48 hours in RPMI medium containing 0–300 mg/L L-glutamine (Gln).

**Supplementary figure S8:**
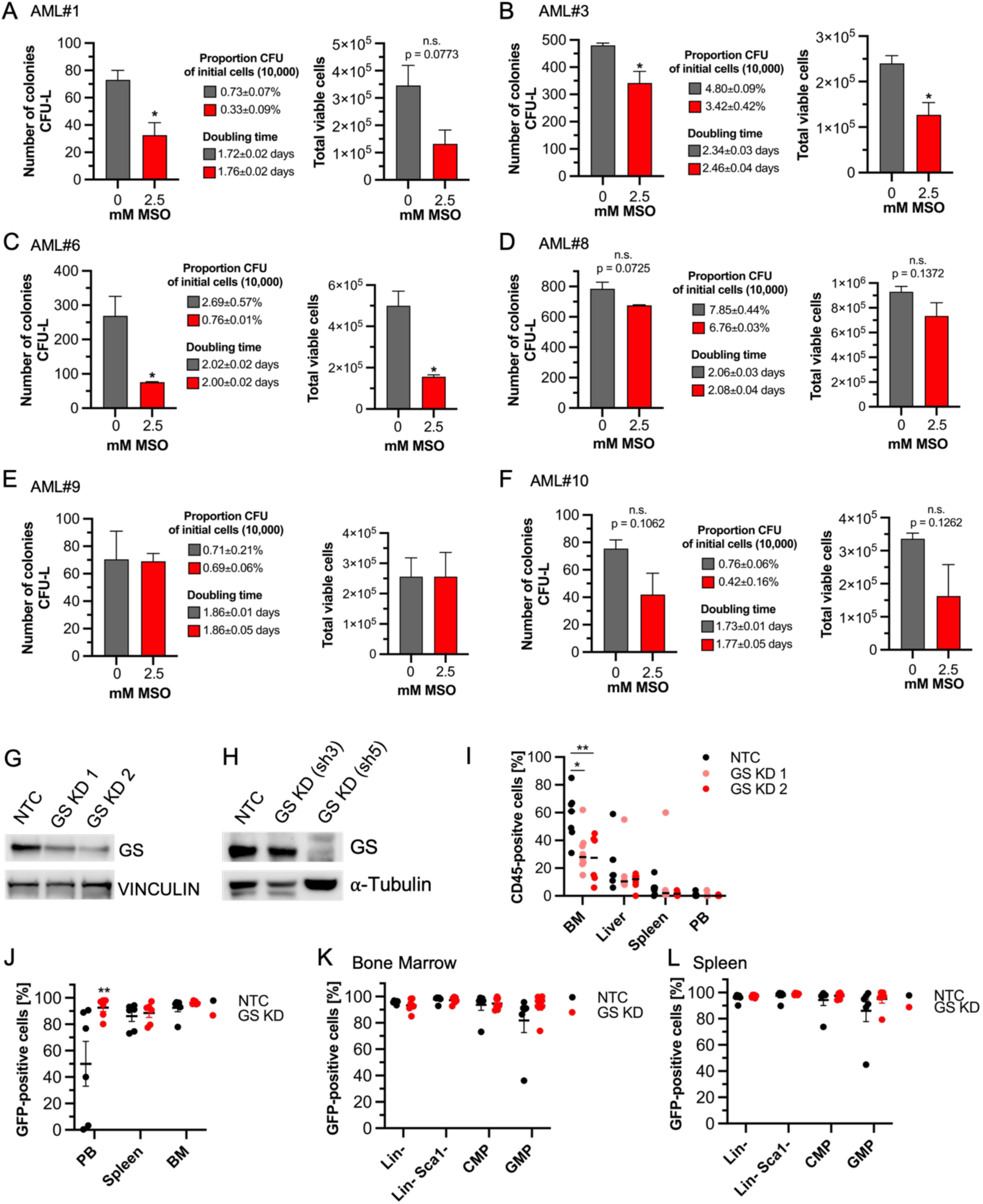
Extended *ex vivo* and *in vivo* evidence for GS dependence in AML. (A–F) Colony formation assay of primary AML blasts treated with 2.5 mM methionine sulfoximine (MSO). Cells were cultured for 21 days under hypoxic conditions. Left: number of CFU-L colonies (bar graphs). Proportion of colony-forming units (CFU) relative to initially seeded cells and doubling time are indicated in the legends. Right: total number of cells in the colony assay. Bar graphs represent mean ± SD of two technical replicates. One-way ANOVA with Bonferroni’s post hoc test; *, p < 0.05. (G, H) Western blot analysis showing GS knockdown efficiency in transplanted cells. Vinculin and α-tubulin served as loading controls. (I) Scatter dot plot showing individual values and the median percentage of CD45-positive cells in bone marrow (BM), liver, spleen, and peripheral blood (PB). Two-way ANOVA with Bonferroni’s post hoc test; *, p < 0.05; **, p < 0.01. (J) GFP-positive cells detected in peripheral blood (PB), spleen, and bone marrow (BM). Only few or very low percentages of GFP-positive cells were detected in the PB of mice that died first. (K, L) GFP-positive cells in bone marrow (K) and spleen (L) within Lin⁻ myeloid progenitors (Lin⁻ CD117⁺ Sca1⁻), common myeloid progenitors (CMP; Lin⁻ CD117⁺ Sca1⁻ CD34⁺ CD16/32⁻), and granulocyte–monocyte progenitors (GMP; Lin⁻ CD117⁺ Sca1⁻ CD34⁺ CD16/32⁺). No GFP-positive cells were detected in megakaryocyte–erythrocyte progenitors (MEP). Scatter dot plots show individual values and mean ± SD. Statistical analysis was performed using Student’s t-test; no significant differences were detected.

## Resource availability

### Lead contact

Information and requests for resources and reagents should be directed and will be fulfilled by the lead contact, Nina Kurrle

### Materials availability

The plasmid pLenti mCherry-TFEB Hygro generated in this study has been deposited to Addgene (252409).

### Data and code availability

- This paper analyzes existing, publicly available data, accessible at Proteomics IDEntifications Database (PRIDE) PXD030463 [39], PXD023201 and PXD028007 [41] and MassIVE repository MSV00089012 [40], and at Bloodspot (servers.binf.ku.dk/bloodspot/): MILE study (GSE13159) [37, 38] and human normal hematopoiesis (GSE42519). Additional information can be found in Supplementary table S5.
- The global proteomics data upon BSA-treatment are accessible at MAssIVE repository (https://doi.org/doi:10.25345/C5PV6BM4R (BSA_Glc)) and are publicly available as of the date of publication and Supplementary table S2
- De-identified patient data can be found in Supplementary table S1
- Global metabolomics data measured by Metabolon is available at National Metabolomics Data Repository (NMDR) (Study ID: ST004761; DOI: http://dx.doi.org/10.21228/M8SK2N) and in Supplementary table S3.
- GSMS data of intracellular tracking of ^15^NH_4_Cl is available at National Metabolomics Data Repository (NMDR) (Study ID ST004845; DOI: http://dx.doi.org/10.21228/M88C39) and in Supplementary table S4.
- Spectra of ^1^H-^15^N heteronuclear NMR experiments are available at National Metabolomics Data Repository (NMDR) (Study ID ST004844; DOI: http://dx.doi.org/10.21228/M8D28B) and are publicly available as of the date of publication.
- Original Western blots are available in Supplementary file S6.
- Any additional information required to reanalyze the data reported in this paper is available from the lead contact upon request

## Supporting information

Supplementary file S6_original Western blots

## Acknowledgments

We thank Jenny Bleeck and Sandra Tzschentke as well as Martine Pape and Marion Bodach (DKTK proteomics platform Frankfurt) for outstanding technical support. Furthermore, we thank the company Silantes GmbH (Munich, Germany) for providing the Arthrospira maxima hydrolysate, Dr. Anjali Cremer (University hospital Frankfurt, Germany) for providing T-ALL and B-ALL cell lines as well es Prof. Dr. Ritva Tikkanen (University of Giessen, Germany) for providing a TFEB WT plasmid. We would also like to thank our colleagues in the wards for their support with the ammonia measurements, in particular Hermine Kunz, Ward B11 and our outpatient clinic, as well as all our patients. We would also like to thank Donatella Novelli, Hanne Platzer, Elke Siebert, Lisa Kuszynski-Bischof, Tessa Schmachtel, Kristina Götze and Birgit Rosiejak for their ongoing organizational support.

## Funding

This research was supported in part by the LOEWE Center Frankfurt Cancer Institute (FCI) funded by the Hessian Ministry of Science and Research, Arts and Culture [III L 5 - 519/03/03.001].

This work was also supported by a grant from the EU (ITN-EJD HaemMetabolome (H2020-MSCA-ITN-2015 675790) awarded to HS, HaS, JS and to MC. JK and IA gratefully acknowledge receipt of a Marie Curie Fellowship and were participants in the same Initial Training Network. HS and NK also acknowledge the support of the Association for Bone Marrow Transplantation and Gene Therapy Frankfurt Main (KGF), a private philanthropic charity formed by patients and friends of the Frankfurt bone marrow transplantation unit.

## Author contributions

Conceptualization, N.K., F.S. and H.S.; Investigation, N.K., P.M. V.S., J.K., I.A. M.S. S.K., C.A.M., S.M, D.F., C.F., N.P., A.B.S., L.F. L.B., H.N., V.St., R.K, M.T., C.M., J.M.; Formal analysis and software, N.K., P.M., S.M., B.H., S.W., J.J., J.J.S., S.S., F.S.; Resources, P.M., V.S., F.G., D.S.K, H.B., S.S., M.C., T.O., H.Sch., H.S.; Supervision, C.B., M.L. T.B., B.B., M.C., T.O., H.Sch., F.S., H.S.; Visualization, N.K. P.M. V.S., J.K., I.A.; Writing-Original draft, N.K., F.S., H.S., All authors read and agreed with the final version of the manuscript.

## Declaration of Interests

T.O. served on advisory boards for and/or received honoraria from AbbVie, BeiGene, Bristol Myers Squibb, Gilead, Janssen, Kite, Kronos Bio, Lilly, Roche, and Sobi. T.O. received travel support from BeiGene, Janssen, Kite, and Roche.

All other authors declare no competing interest related to this work.

## Declaration of generative AI and AI-assisted technologies in the writing process

During the preparation of this work the authors used ChatGPT (OpenAI) in order to assist with code development for data analysis in Jupyter notebooks and to refine scientific text for clarity and style. After using this tool, the authors reviewed, verified and edited all generated matters and take full responsibility for accuracy and integrity of the final manuscript.

