## Supplementary file S6_original Western blots for "Extracellular protein catabolism drives regulated nitrogen handling and ammonia buffering in acute myeloid leukemia"

**Figure 2: Extracellular protein catabolism sustains energy metabolism and engages nitrogen-handling pathways in AML.**

**(D):**

Phospho-P70S6K (T389):

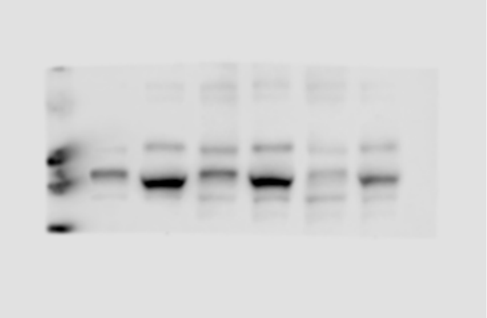

Vinculin of phospho-P70S6K blot:

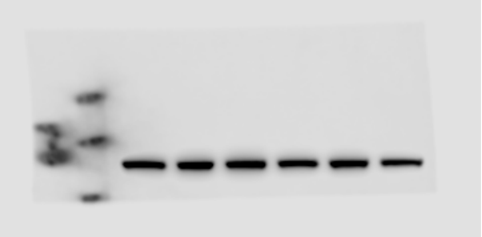

P70S6K:

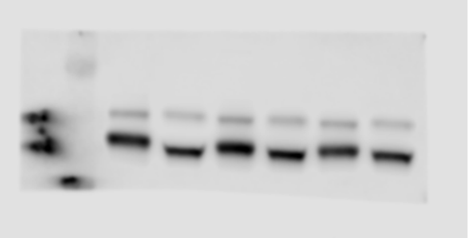

Vinculin of P70S6K blot:

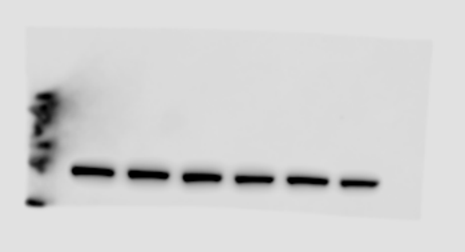

Phospho-4EBP1(S65):

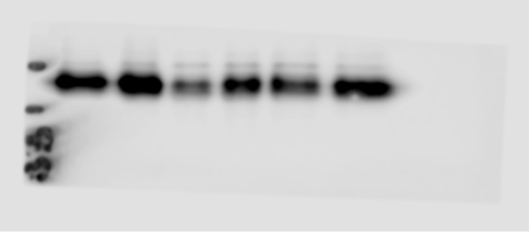

Vinculin of phospho-4EBP1 (long and short exposure)

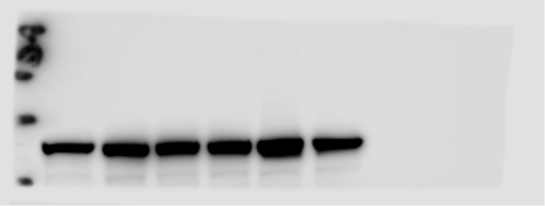

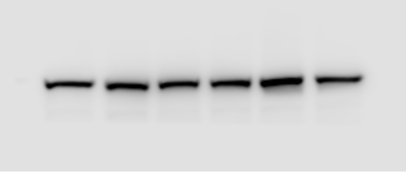

4EBP1:

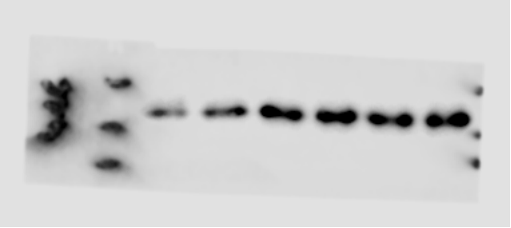

Vinculin of 4EBP1:

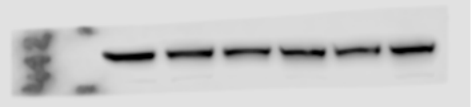

**Figure 4: Glutamate–ammonia ligase links ammonia assimilation to extracellular protein catabolism in AML.**

**(D):**

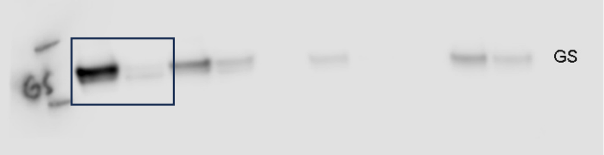


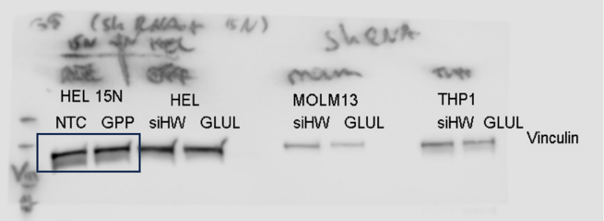

**Supplementary figure S5: Extracellular albumin supplementation activates anabolic signaling and drives broad metabolic remodeling in AML cells.**

**(B)**

SREBP2:

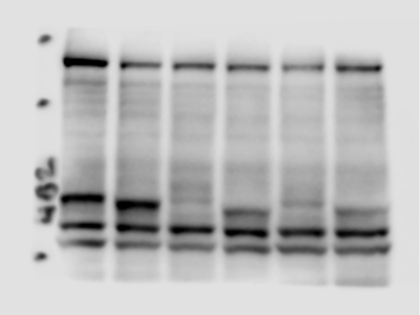

SQLE:

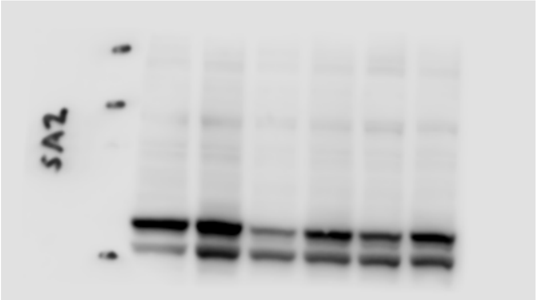

Histone H3 of SREBP2:

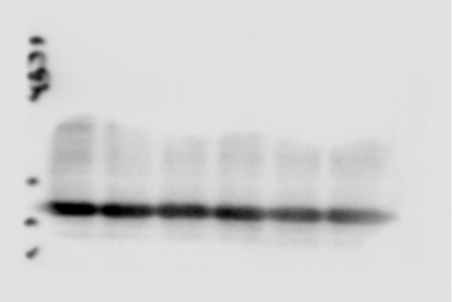

Histone H3 of SQLE:

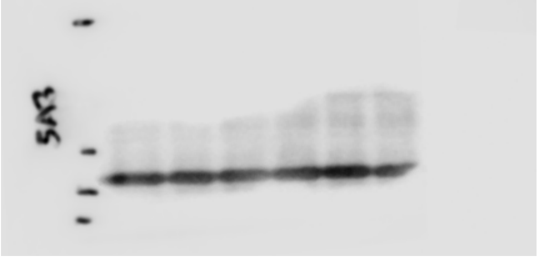

**(C)**

Phospho-TFEB (S211):

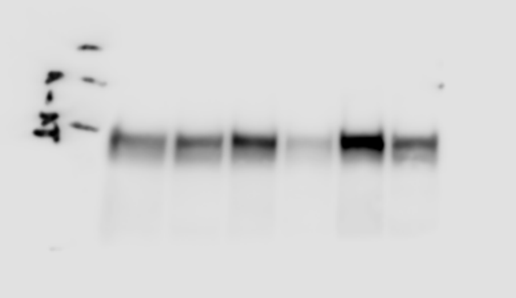

TFEB total:

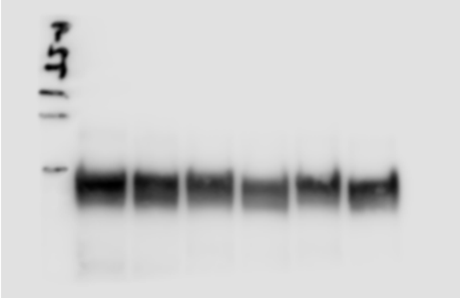

Vinculin of Phospho-TFEB S211 (right) and TFEB total (left):

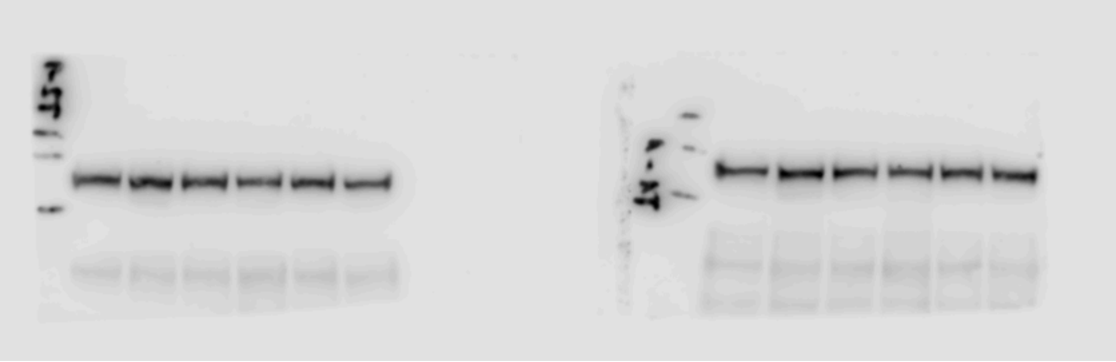

Phospho-TFEB S122:

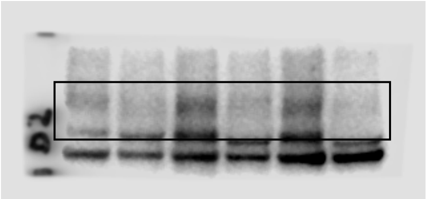

Vinculin of phospho-TFEB S122:

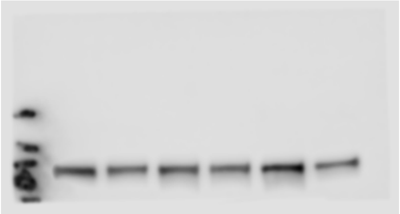

**(D)**

Phospho-TFEB (S211): (short and long exposure)

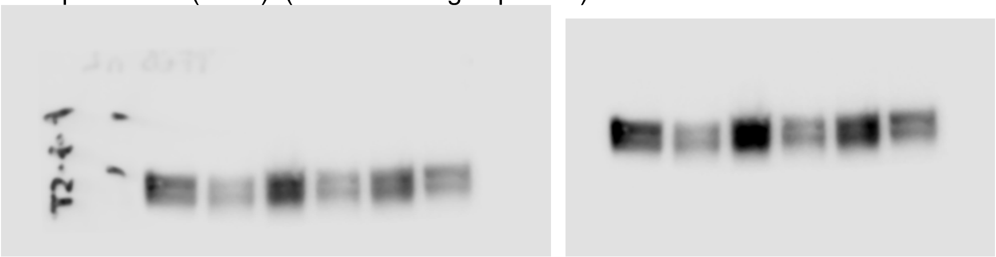

TFEB:

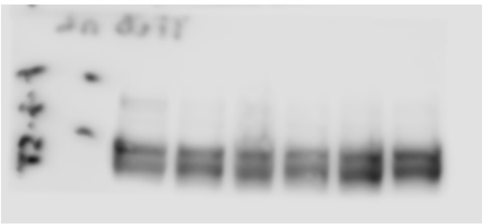

Beta-Tubulin: (short and long exposure)

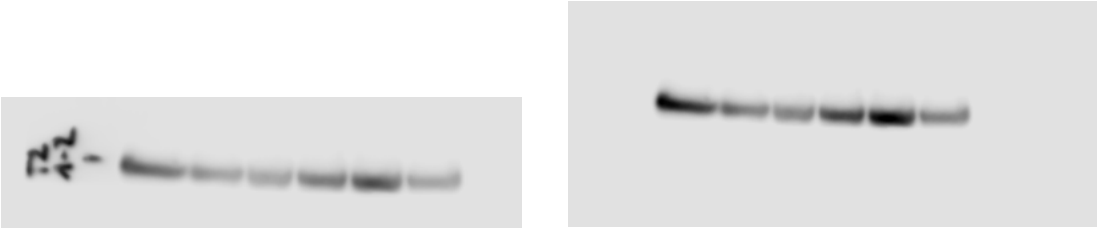

Phospho-TFEB (S122): (short and long exposure

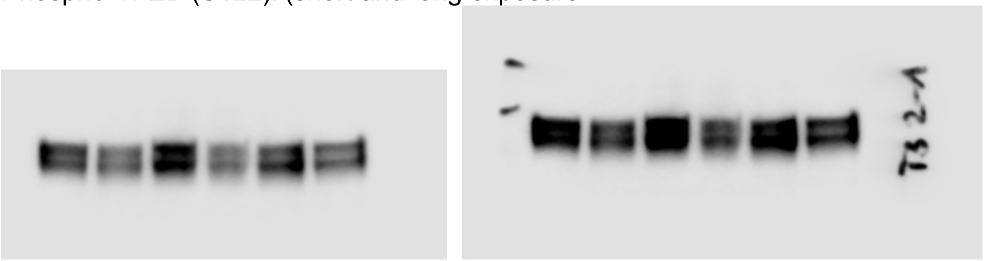

Beta-Tubulin: (short and long exposure)

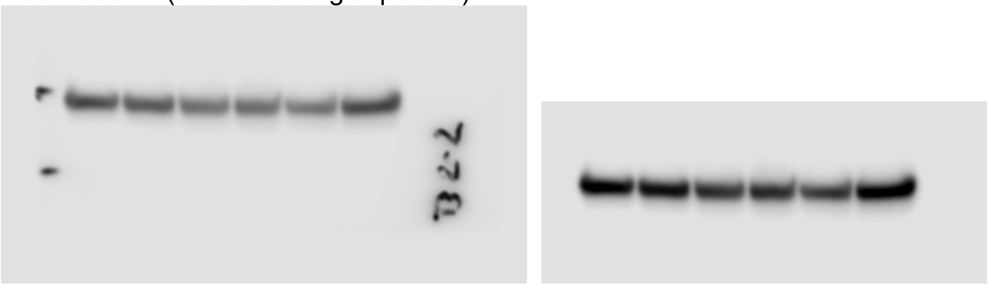

**Supplementary figure S6: Bone marrow ammonia distributions and supporting data for ammonia assimilation and tracing experiments**

**(C)**

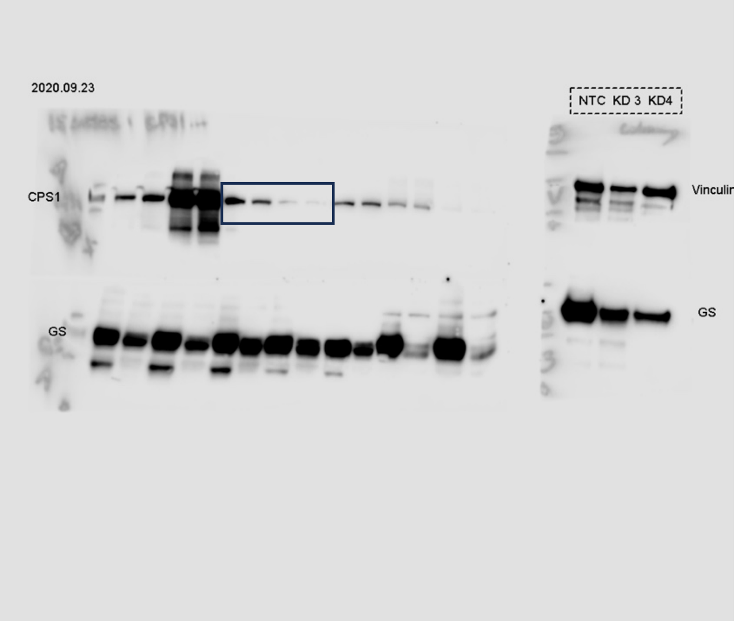


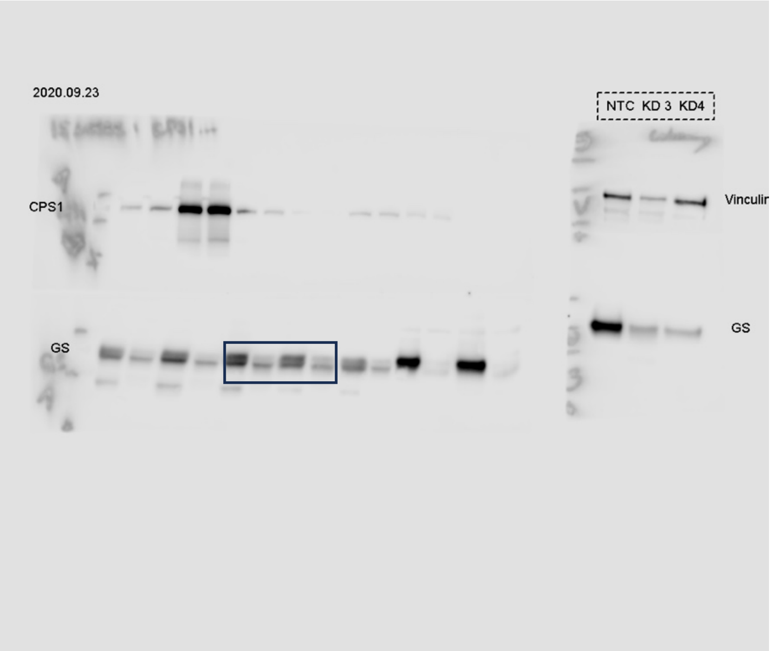

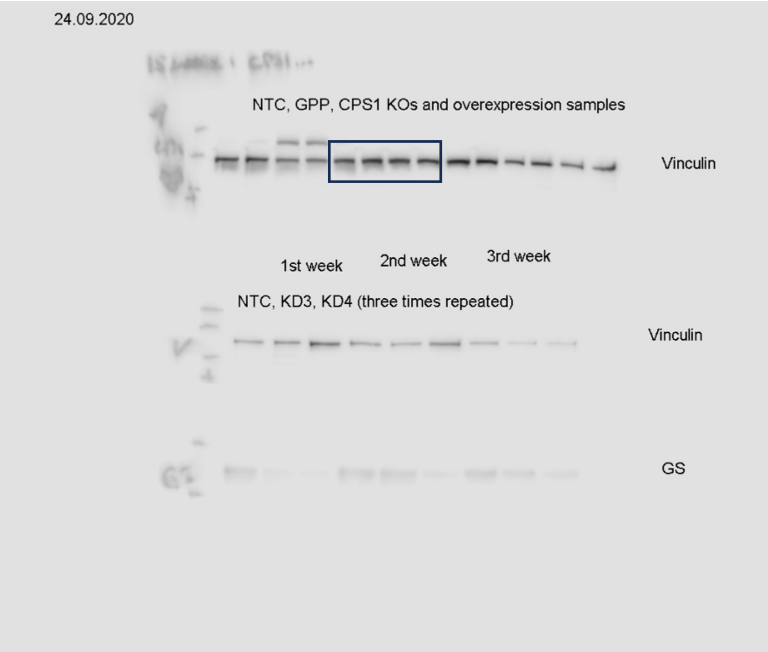


**Supplementary figure S7: GS expression across cancer models and validation of GS genetic pertubation in AML cells.**

**(E)**
GS (lower band) and Nucleoline (upper band):

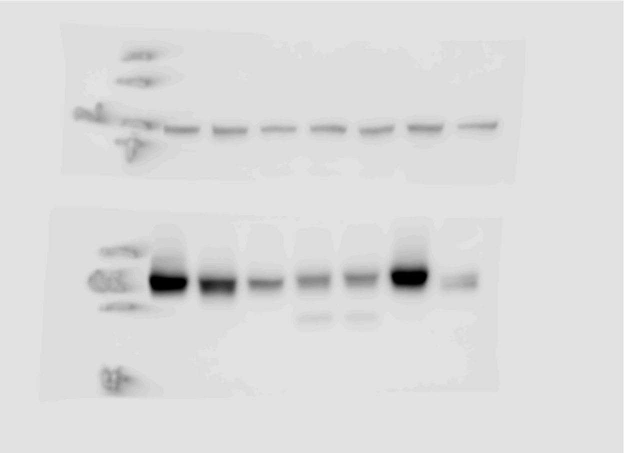

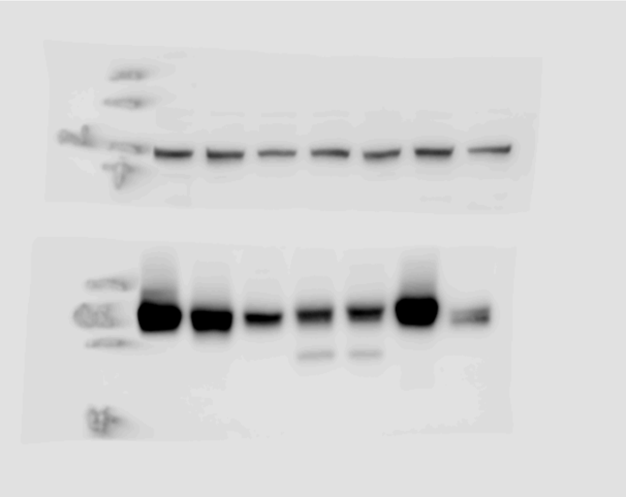

**(I)**

GS:

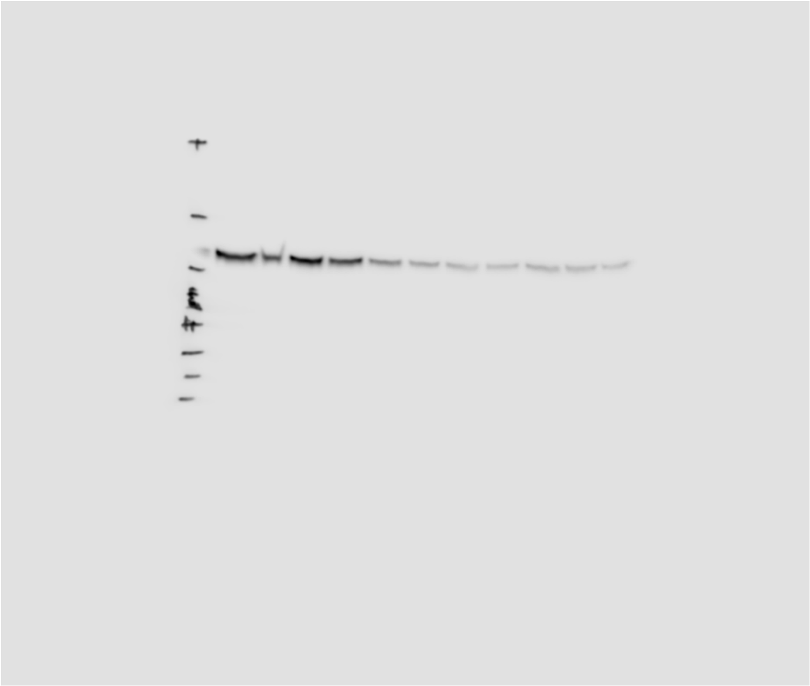

Vinculin:

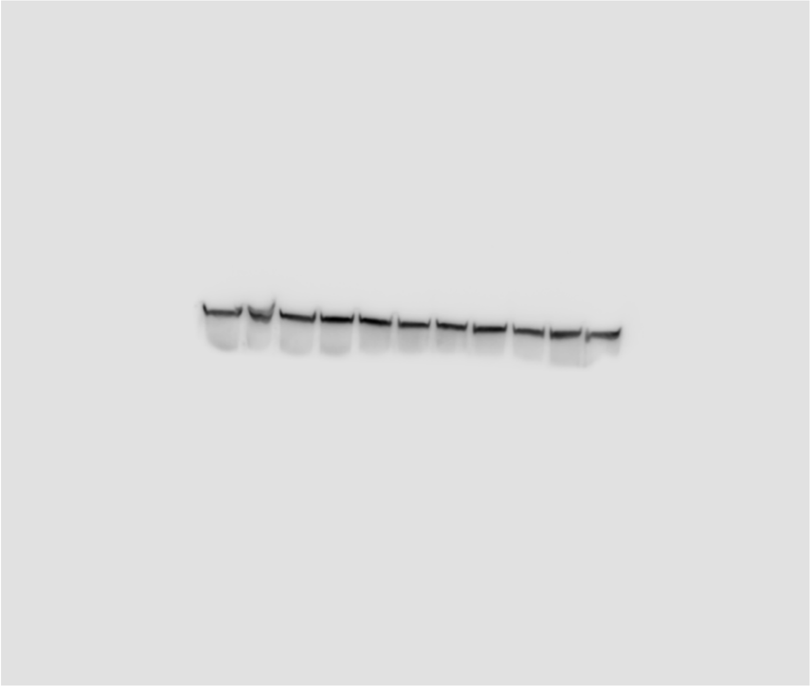

**Supplementary figure S8: Extended *ex vivo* and *in vivo* evidence for GS dependence in AML.**

**(G)**

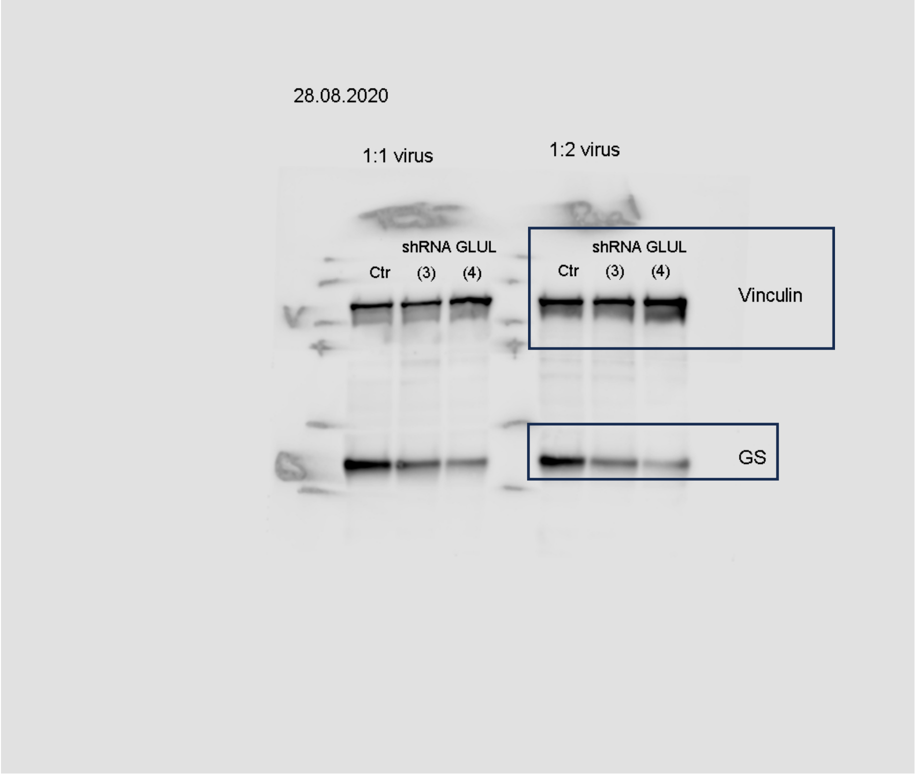


**(H)**

GS:

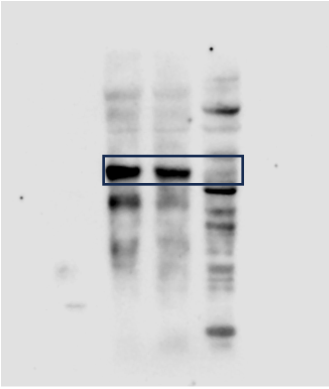

Alpha-tubulin

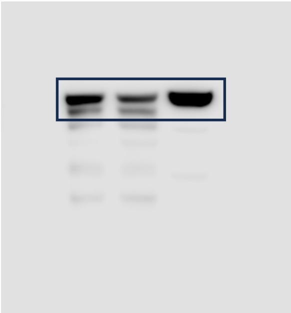
